# Brainstem GLP-1 neurons modulate physiological satiation and drive sustained weight loss in obese mice

**DOI:** 10.64898/2025.11.29.691273

**Authors:** Wanqing Jiang, Cecilia Skoug, Ian Rodrigues, Ernesto Ciabatti, Fiona M. Gribble, Frank Reimann, Daniel I. Brierley, Marie K. Holt, Stefan Trapp

## Abstract

Glucagon-like peptide-1 receptor (GLP-1R) activation in the brain strongly reduces appetite, but most brain GLP-1Rs are not accessible for systemically administered GLP-1R agonists. Acute activation of nucleus tractus solitarius (NTS) GLP-1 neurons, targeting brain GLP-1Rs, strongly suppresses food intake separate from GLP-1R agonists. However, it is unknown if their chronic stimulation is a viable strategy for appetite suppression, or if obesity disrupts their function. Here we demonstrate that GLP-1 neurons, distributed through NTS and IRT, determine meal size and their number is inversely correlated with bodyweight gain. GLP-1 neurons in IRT and NTS differ in their inputs but generate outputs in largely overlapping areas. Higher body weight and fat mass predicts a higher percentage of active GLP-1 neurons. Their chemogenetic activation is an efficient means of reducing food intake, and in obese mice chronic activation elicits sustained weight loss. In conclusion, GLP-1 neurons are a feasible target for obesity treatment.

## Introduction

Glucagon-like peptide-1 (GLP-1) receptor agonists (GLP-1RAs), such as semaglutide^1^, have transformed obesity treatment, eliciting >15% weight reduction in overweight adults after >1year of weekly dosage^2, 3^. This weight loss is primarily driven by a reduction in food intake, caused by GLP-1RAs acting on the limited subset of CNS GLP-1 receptors which are accessible from the systemic circulation^4–9^, mimicking the action of supraphysiological concentrations of gut-derived peripheral GLP-1. However, GLP-1 is also produced by neurons in the brain, and this central GLP-1 system can supress food intake independently of the peripheral GLP-1 system, and is not recruited by GLP-1RA drugs for their anorectic effects^10–12^. Stimulating endogenous brain GLP-1 signalling may thus represent a valuable novel target for obesity pharmacotherapy, with unrealised potential to increase the efficacy and/or tolerability of current GLP-1RAs. To realise this potential, however, crucial knowledge gaps regarding the roles of endogenous central GLP-1 signalling in eating and bodyweight control must be addressed.

Within the brain, GLP-1 is primarily produced by brainstem preproglucagon (PPG) neurons, also known as *Gcg* neurons^13, 14^. Their cell bodies are located in the caudal nucleus tractus solitarius (NTS; hence PPG^NTS^ neurons) and the intermediate reticular nucleus (PPG^IRT^ neurons), from where they project widely throughout the CNS, but not to the periphery^13, 15–20^. Numerous studies have now shown that acute chemogenetic or optogenetic activation of PPG^NTS^ neurons robustly suppresses eating in lean mice, without evidence of negative valence or behavioural disruption^12, 14, 21, 22^. However, loss-of-function manipulations of this NTS population have provided limited evidence for their role in physiological meal control under *ad libitum* feeding conditions^13^. This apparent discrepancy suggests that the IRT population may have a role in physiological meal control, but studies to date have largely ignored this population.

Notably, previous studies have shown that ablation of PPG^NTS^ neurons leaves approximately 30% of GLP-1 tissue content remaining in the brainstem and 20% in hypothalamus^13^ which likely originates from PPG^IRT^ neurons. This leaves open the possibility that this remaining GLP-1 provides a crucial but unappreciated contribution to physiological meal control, not captured by loss of function studies to date focussed on the NTS population alone. Similarly, the translational potential of targeting central GLP-1 release for weight loss may have been underestimated by activation studies which have omitted the IRT population, which represents ∼35% of all brainstem PPG neurons. It is also unclear whether the NTS and IRT populations are effectively a continuous population with consistent connectivity patterns, or the extent to which their afferent and efferent connectivity converges and/or diverges, and the functional implications thereof.

Here we investigated these two brainstem PPG neuron populations, individually and in combination, to compare their circuit connectivity, roles in physiological appetite regulation, and their translational potential based on capacity to suppress eating in a sustained manner in obese mice. We report that PPG^NTS^ and PPG^IRT^ neurons receive distinct monosynaptic inputs, but have convergent efferent projection targets throughout the brain, and that combined ablation of both populations delays the onset of physiological satiation to a degree sufficient to promote weight gain under *ad libitum* chow fed conditions. Crucially, chronic daily chemogenetic activation of PPG^NTS+IRT^ neurons drives robust and sustained hypophagia and weight loss in obese mice without notable adverse effects, demonstrating their value as targets for obesity pharmacotherapy.

## Results

### PPG^NTS^ and PPG^IRT^ neurons have distinct inputs but convergent projections

It is currently unclear whether PPG^NTS^ and PPG^IRT^ neurons comprise a homogenous population with parallel circuit architecture, or if they have distinct afferent and/or efferent connectivity which may underlie functional specialisation within the central GLP-1 system. We thus first addressed whether PPG^NTS^ and PPG^IRT^ neurons have distinct projection patterns using adeno-associated virus (AAV)-mediated selective expression of tdTomato in either of these populations in adult PPG-Cre-YFP mice (Figure 1a,b). This enabled specific labelling of PPG^NTS^ or PPG^IRT^ neurons and their respective axonal projections (tdTomato^+^). Colocalization of YFP fluorescence, representing the entire PPG neuron population, in tdTomato-expressing axons validated the specificity of this approach (Suppl Fig 1). PPG^IRT^ neurons constituted 31±4% of the total brainstem PPG neuron population (based on YFP fluorescence). We successfully targeted 84±5% PPG^NTS^ and 69±7% PPG^IRT^ cell bodies (based on tdTomato fluorescence; Fig 1a,b). Axon projections from PPG^NTS^ and PPG^IRT^ neurons were found in largely overlapping regions extending from the brainstem to the forebrain (Figure 1c,d, Suppl Figure 1).

**Figure 1.**
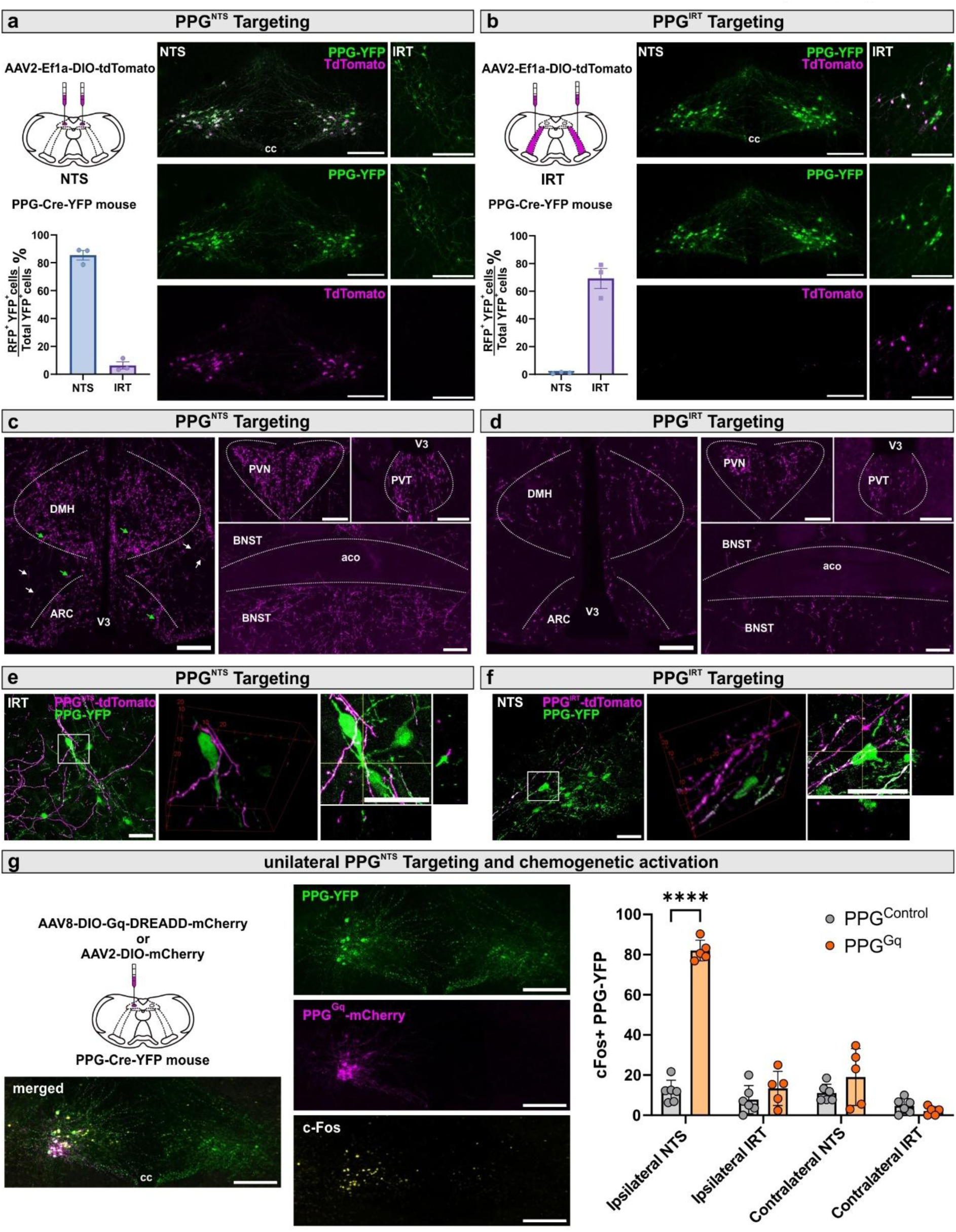
PPG^NTS^ and PPG^IRT^ neurons have overlapping projection patterns but do not form a highly interconnected network. **a,b**. Schematic of bilateral viral injection into both NTS or IRT regions to induce tdTomato fluorescence expression in the PPG^NTS^ or PPG^IRT^ neurons (n=3, each) of PPG-Cre-YFP mice. Photomicrographs showing expression of tdTomato fluorescence in YFP expressing PPG neurons. Scalebars: 200μm. cc, central canal. Quantification of tdTomato immunoreactive YFP^+^ PPG neurons in the NTS and IRT. **c,d.** tdTomato-immunoreactive axons originated from PPG^NTS^ or PPG^IRT^ neurons. DMH, dorsal medial hypothalamus; ARC, arcuate nucleus; PVN, paraventricular nucleus, PVT: paraventricular thalamus; BNST, bed nucleus of the stria terminalis. Green arrows: varicose axons; v3: third ventricle; aco: anterior commissure. White arrows: smooth axons of passage. Green arrows: varicose axons. Scale bars: 200μm. **e,f.** Maximum projection of confocal image z-stack showing tdTomato-immunoreactive axons originated from PPG^NTS^ or PPG^IRT^ neurons and YFP^+^ PPG^NTS^ or PPG^IRT^ neuron cell bodies and their dendrites. Rotated 3D view and orthogonal views depicting an YFP^+^ PPG neuron with close apposition of tdTomato-immunoreactive axons originating from the other PPG neuron population. Scale bars: 20 μm. **g.** Schematic of viral injection and experimental paradigm for unilateral chemogenetic activation of PPG^NTS^ neurons. Photomicrograph of cFos immunoreactivity (cFos-ir) in PPG^NTS^ and PPG^IRT^ neurons. Scale bars: 200μm. cc, central canal. Quantification of cFos-ir+ PPG^NTS^ and PPG^IRT^ neurons in control and ChR2 mice, 2-way ANOVA, region x virus F (1, 10) = 27.55, ***p=0.0004, NTS: **** p<0.0001. All data are presented as mean±SEM.

Notably, occasional axon varicosities from both PPG^NTS^ neurons and PPG^IRT^ neurons were found in close proximity to PPG cell bodies and processes of the respective other region (Figure 1e,f). In view of a previous report of numerous close appositions between PPG neurons within the NTS^23^, we investigated whether functional connectivity exists between contralateral PPG^NTS^ neurons and between PPG^NTS^ neurons and ipsi- and/or contralateral PPG^IRT^ neurons, using cFos immunoreactivity induced by unilateral chemogenetic activation of PPG^NTS^ neurons (Fig 1g). Whilst the hM3Dq-transduced PPG^NTS^ neurons themselves exhibited strong cFos immunoreactivity, no increase in cFos beyond baseline levels was observed in contralateral PPG^NTS^ neurons nor in PPG^IRT^ neurons on either side of the brainstem. In contrast, a significant increase in cFos immunoreactivity was observed in projection targets from the PPG neurons, such as the PVN (suppl Fig 2a). Similarly, optogenetic stimulation of PPG^NTS^ neurons failed to induce cFos immunoreactivity in PPG^IRT^ neurons (suppl Fig 2b). Altogether, these data demonstrate that PPG^NTS^ and PPG^IRT^ neurons have a largely converging pattern of efferent connectivity but provide no evidence for direct functional connectivity from PPG^NTS^ to PPG^IRT^ neurons. This suggests these two PPG neuron populations do not constitute a sequential feedforward circuit, but provide largely parallel, disparate inputs that converge on multiple CNS target regions with well-established roles in eating and bodyweight regulation.

To map and compare the afferent inputs to each PPG neuron population, we conducted monosynaptic retrograde rabies tracing in PPG-Cre-YFP mice^24, 25^. This approach enabled us to identify neurons that provide monosynaptic inputs to PPG neurons targeted by the mCherry-encoding rabies virus (Fig 2a,b).

**Figure 2.**
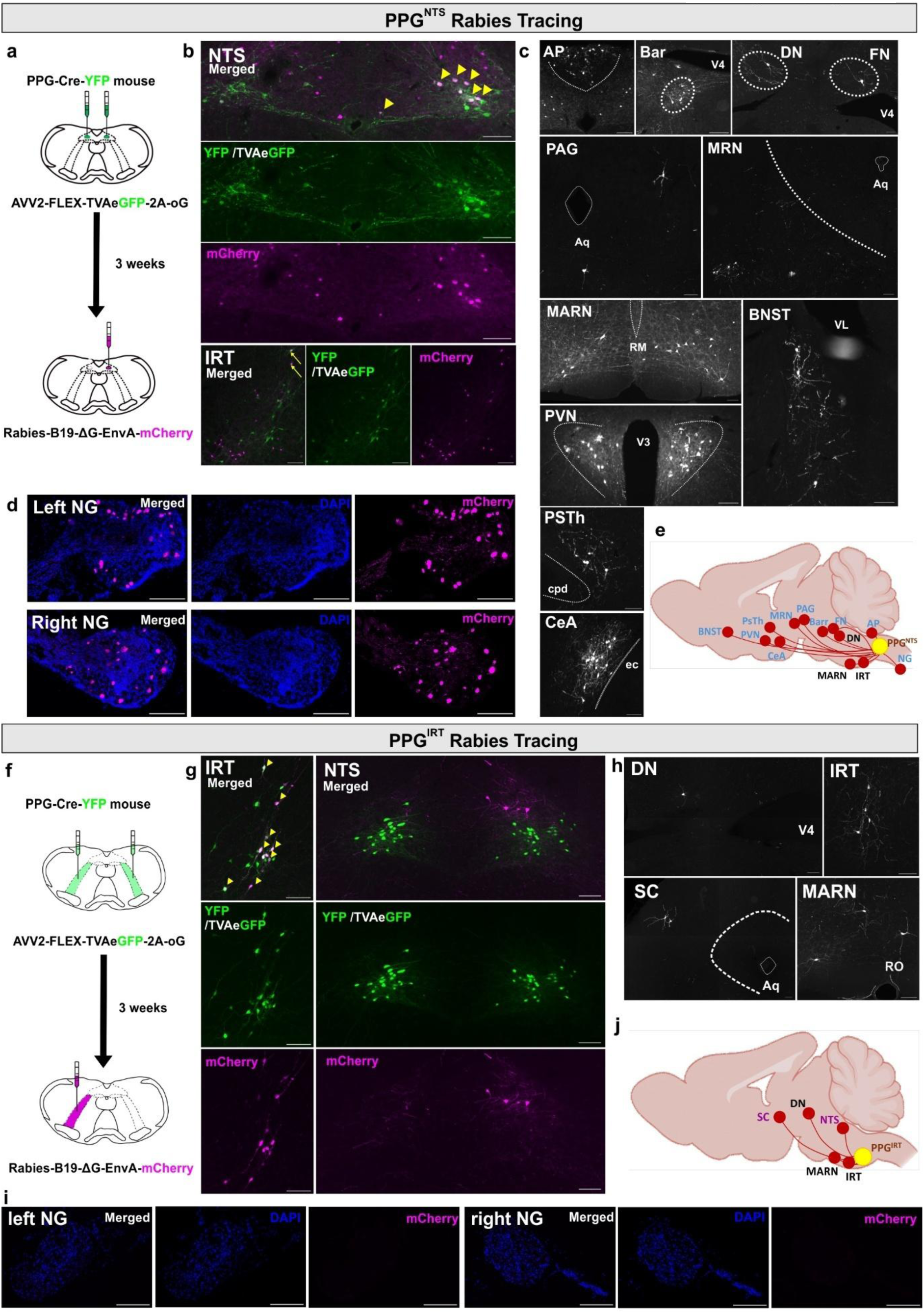
PPG^NTS^ and PPG^IRT^ neurons receive distinct monosynaptic inputs and vagal afferent neurons only provide input to PPG^NTS^ neurons. a,f Schematics illustrating the viral injection in PPG-Cre-YFP mice for unilateral monosynaptic retrograde rabies viral tracing of either PPG^NTS^ (**a**, n=2) or PPG^IRT^ neurons (**f**, n=3). **b,g** Photomicrographs showing the NTS (**b**) or IRT (**g**) regions that received viral injections, with ’starter’ neurons indicated by yellow arrowheads, co-expressing Rabies mCherry (detected with anti-dsRed antibody) and TVA-GFP or mouse YFP (detected with anti-GFP antibody). Cells which serve as input sources are detected by Rabies-mCherry immunofluorescence labelling. Yellow arrows in the lower panel of (b) indicate PPG^IRT^ neurons, located on the opposite side of the viral injection, as input sources to PPG^NTS^ neurons and co-express both Rabies-mCherry and TVA-GFP or mouse YFP. **c,h** Additional central and peripheral source of inputs to the PPG^NTS^ (**c**) and PPG^IRT^ (**h**) neurons. **d,i** Detection of cells in both the left and right nodose ganglia (NG) as source of inputs to PPG^NTS^ (**d**) but not PPG^IRT^ (**i**) neurons. **e,j** Schematics summarising the source of input to PPG^NTS^ (**e**) and PPG^IRT^ (**j**) neurons. For PPG^IRT^ neurons, the schematic includes only regions that were consistently identified across all three mice in the experiment. The shared source of inputs to both PPG population are indicated in black. Abbreviations: DN, dentate nucleus; SC, superior colliculus; MARN, magnocellular reticular nucleus; AP, area postrema; Barr, Barrington’s nucleus; FN, fastigial nucleus; PAG, periaqueductal gray; MRN, midbrain reticular nucleus; BNST, bed nucleus of the stria terminalis; PVN, paraventricular nucleus; PSTh, parasubthalamic nucleus; CeA, central amygdala; cc, central canal; V4, fourth ventricle; Aq, cerebral aqueduct; RO, nucleus raphe obscurus; RM, nucleus raphe magnus; VL, ventral lateral; V3, third ventricle; cpd, cerebral peduncle; ec, external capsule. Scale bars: 100μm.

As reported previously^26^, PPG^NTS^ neurons receive inputs from widespread subcortical regions. In the telencephalon these include the BNST and CeA, whilst in the hypothalamus PVN and PSTh were found to provide dense input (Fig 2c,e). PPG^NTS^ neurons also receive input from midbrain and pontine regions, including the reticular nucleus (MRN), periaqueductal grey (PAG) and Barrington’s nucleus (Bar). We also identified inputs from the cerebellar dentate and fastigial nucleus (DN and FN), as well as areas of the medulla, including the magnocellular reticular nucleus (MARN), contralateral IRT and the area postrema (AP). Finally, PPG^NTS^ neurons receive peripheral sensory inputs via the afferent vagal nerve (Fig 2d). Interestingly, PPG^NTS^ neurons were found to receive inputs from a very few PPG^IRT^ neurons (Fig 2b).

In contrast, only a few brain regions were shown to provide monosynaptic input to PPG^IRT^ neurons and these were restricted to pontine, cerebellar and medullary regions, with no clear input from the telencephalon or hypothalamus (Fig 2 f-j). Comparison of the inputs to PPG^IRT^ and PPG^NTS^ neurons revealed shared inputs from the DN, MARN and the contralateral IRT. PPG^IRT^ neurons receive additional inputs from the NTS, as well as the superior colliculus (SC). In contrast to PPG^NTS^ neurons, the IRT population was devoid of vagal afferent input.

In summary, our anterograde and retrograde tracing and functional connectivity studies demonstrate that the PPG^NTS^ and PPG^IRT^ neuron populations each integrate multiple and largely distinct inputs and convey these integrated signals in a divergent but parallel manner to overlapping projection target regions. This implies that the convergent inputs of these populations are integrated by their postsynaptic targets to induce functional effects on eating and bodyweight. Previous studies targeting PPG^NTS^ neurons alone may have thus underestimated both the physiological relevance of the overall PPG population, and its translational potential for inducing hypophagia-driven weight loss. We thus sought to test this hypothesis by conducting loss-of-function and activation manipulations targeting the brainstem PPG neuron population in its entirety.

### Combined ablation of PPG^NTS^ and PPG^IRT^ neurons impairs satiation and increases body weight

Brainstem PPG neurons were ablated in adult PPG-Cre-YFP mice of either sex using AAV-mediated selective expression of diphtheria toxin A subunit (DTA; Fig 3a-c). Quantitative analysis in PPG^DTA^ mice and PPG^Control^ mice at the end of the study revealed that all PPG^NTS^ neurons were ablated and the majority of the PPG^IRT^ neurons (Fig 3d,h), resulting in an almost complete loss of PPG axons in projection targets such as the PVN (suppl Fig 3a).

**Figure 3.**
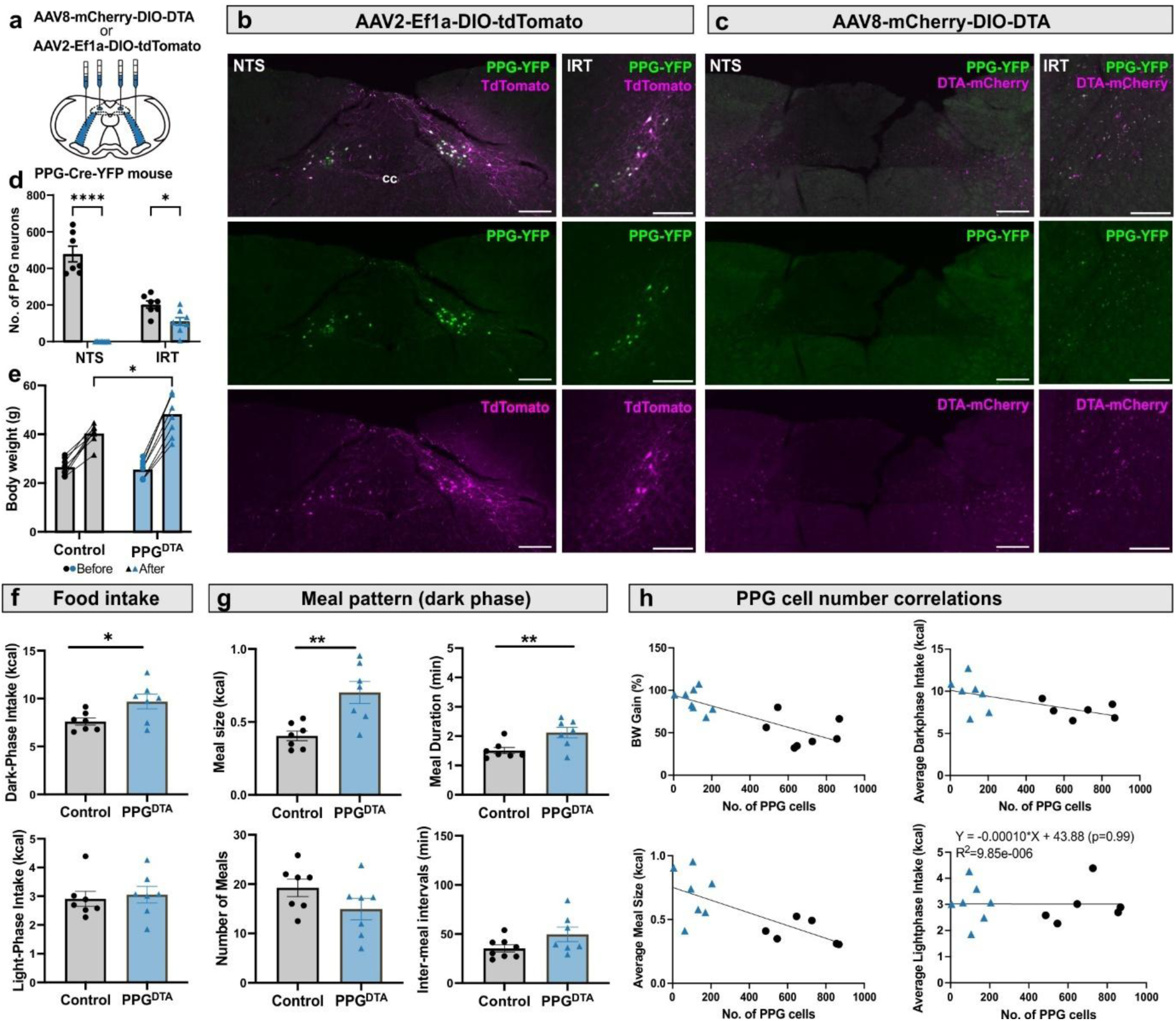
Ablation of PPG^NTS^ and PPG^IRT^ neurons significantly increases body weight and *ad libitum* food intake with a correlation between these parameters and the number of functional PPG neurons. PPG-Cre-YFP mice received bilateral viral injections into both NTS and IRT for either PPG neuron ablation (PPG^DTA^, n=8) or tdTomato fluorescence expression as controls (PPG^Control^, n=8). **a** Schematic illustrating bilateral viral injections. **b,c** Photomicrographs showing specific ablation or tdTomato fluorescence expression in PPG neurons in the NTS and IRT. mCherry is expressed in a Cre-independent manner and marks the area of viral transduction. Scale bars: 200 μm. cc: central canal. **d** Quantification of total YFP^+^ PPG neurons in PPG^Control^ (grey bars) and PPG^DTA^ (blue bars) mice. Data are presented as mean ± SEM. Two-way ANOVA, region × virus: F(1, 26) =59.61, ****p<0.0001. NTS: ****p<0.0001, IRT: *p=0.03. **e** Body weight of PPG^Control^ and PPG^DTA^ mice before and 6 months after surgery. Two-way ANOVA, time × virus: F(1, 14) = 14.77, **p=0.002; Sidak post hoc test, after surgery: *p=0.012. **f** PPG neuron ablation increased dark-phase food intake without affecting light-phase food intake. Data are averaged across 6 experimental days and presented as mean ± SEM. Unpaired nested t-test, dark-phase food intake: *p=0.03. **g** PPG neuron ablation increased satiation. Data are averaged across 6 experimental days and presented as mean ± SEM. Unpaired nested t-test, meal size: **p=0.004, mean meal duration: *p=0.01. **h** The number of YFP^+^ PPG neurons was negatively correlated with percentage body weight gain (slope = -0.063 ± 0.013, p = 0.0004, R^2^ = 0.64), meal size (slope = -0.0072 ± 0.0021, p = 0.005, R^2^ = 0.53), and 12h dark-phase food intake (slope = -0.050 ± 0.021, p = 0.03, R^2^ = 0.36), but not with 12h light-phase intake. Linear regression analysis.

Mice of both treatment groups had similar body weights before surgery, but body weight gain differed post-surgery with the PPG^DTA^ group becoming significantly heavier compared to the PPG^Control^ group (Fig 3e). PPG^DTA^ mice ate significantly more chow during the dark, but not light, phase compared to PPG^Control^ mice (Fig 3f). Meal pattern analysis revealed that this was driven by intake of significantly larger meals with a longer meal duration. There was no significant difference in the number of meals or inter-meal intervals, indicating an impairment of satiation under standard *ad libitum* chow fed conditions, but unchanged satiety in PPG^DTA^ mice (Fig 3g).

A significant negative correlation was found between the number of PPG neurons and percentage body weight gain, dark-phase food intake and meal size, but not light phase intake (Fig 3h). These results indicate that PPG neurons are required for physiological satiation, to the extent that they are necessary for maintenance of normal bodyweight with the number of functional PPG neurons determining the size of the effect. Interestingly, this correlation is strongly driven by the control mice which revealed a considerable individual variability in the number of brainstem PPG neurons. This population variability is further explored in the experiments described in Fig 6d, which quantified variability in cFos activation in PPG neurons between mice.

### Chronic PPG neuron activation transiently reduces food intake and body weight in lean mice only

To assess the potential of PPG neurons as a target for anti-obesity treatment, we next investigated whether repeated activation of both PPG^NTS^ and PPG^IRT^ (PPG^Gq^) neurons using a daily dose of 2 mg/kg CNO over two weeks could produce sustained weight loss (Fig 4a). In lean, chow-fed mice, the first dose of CNO led to a significant reduction in food intake, water intake and energy expenditure (EE), with the main effect occurring within the first 3–4 hours (Suppl Fig 4a), consistent with both the time window of effect reported in previous studies^12^ and the known pharmacokinetics of CNO. No compensatory increase in eating was observed for the rest of the day in PPG^Gq^ mice (Suppl Fig 4a), leading to significantly reduced total food intake (Fig 4b) and 2.8+/-0.8% weight loss on day 1. The reduction in food intake was driven by both increased satiation and satiety, reflected in shorter meal duration and longer inter-meal intervals, as well as fewer meals (Fig 4c).

**Figure 4:**
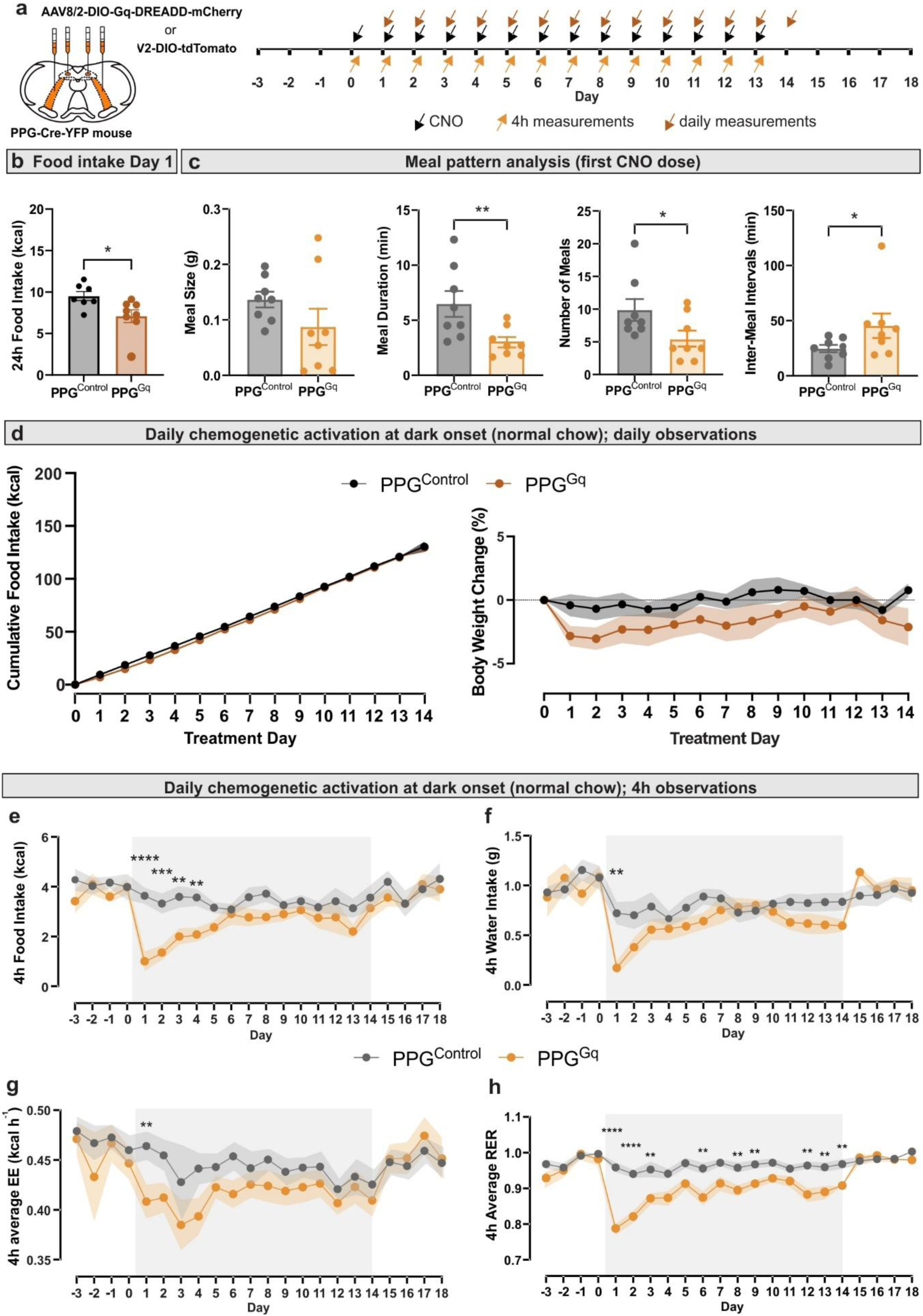
Repeated chemogenetic activation of PPG^NTS^ and PPG^IRT^ neurons only transiently decreases food intake and body weight in the lean state. **a** Schematic representation of bilateral viral injections of AAV2-DIO-tdTomato (PPG^Control^, n=8) or AAV8-DIO-hM3Dq-mCherry (PPG^Gq^, n=8) into the NTS and IRT of PPG-Cre mice and timeline of the experiment once stable viral expression was achieved. Mice were introduced to the Promethion^TM^ cages 3 days prior to the first of 14 daily CNO (2mg/kg) i.p. injections (black arrows) 30 min prior to dark onset. Daily measurements of cumulative food intake and bodyweight (orange arrows) were performed 9h into the next light phase, when the 3h fast prior to dark onset was started. Acute measurements of CNO effects (blue arrows) were taken 4h after dark onset. All 4h measurements and daily food intake measurements were taken from continuous Promethion^TM^ data recording without disturbing the mice. **b** Cumulative food intake for the 24h after the first dose of CNO was significantly reduced in PPG^Gq^ mice. **c** This was due to a significant reduction in meal duration and number of meals as well as an increase in inter-meal interval duration. **d** Cumulative food intake and body weight change were not significantly different between PPG^Gq^ and PPG^control^ mice of 14 days of daily CNO administration. Body weight change (%) was calculated by normalising to the body weight on the day before treatment started (day-1). 2-way ANOVA: Virus: F(1,14) =2.40, p=0.144, Time: F(5.29,74.01)=2.13, p=0.07, Time x Virus: F(13,182)=0.75, p=0.71. **e** Food intake over 4 hours from dark onset for each treatment day. 2-way ANOVA: Time x Virus: F(13,182)=2.39, **p=0.005, Fisher’s LSD test: Day1: ****p<0.0001, Day2: ***p=0.0002, Day3: **p=0.002. Day4: **p=0.004. **f** Water intake over 4 hours each day. 2-Way ANOVA: Time x Virus: F(13, 182) = 2.64, **p=0.002, Fisher’s LSD test: Day1: **p=0.001. **g** Average energy expenditure (EE) over 4h post dark onset each day. Data analysed using ANCOVA with adjustment to body weight for each treatment day with a significance threshold set at p value<0.01. Day1: virus: F (1,13)=11.53, **P=0.005, body weight: F(1,13)=8.78, *p=0.01. **h** Average 4h respiratory exchange ratio (RER): 2-Way ANOVA: Time x virus: F (13,182) = 3.67, **** P<0.0001. Fisher’s LSD test: Day1: ****p<0.0001, Day2: ***p<0.0001, Day3: **p=0.01, Day6: **p=0.008, Day8: **p=0.009, Day9: **p=0.005, Day 12: **p=0.003, Day13: **p=0.01, Day14: **p=0.003. 2-way ANOVA was conducted for the 14 treatment days, followed by Fisher’s LSD post-hoc test with a significance threshold set at p value<0.01. All data are presented as mean±SEM.

Following this, mice continued to receive daily CNO injections for 14 days to achieve repeated activation of PPG neurons. As the primary effects were expected within the first four hours post-injection, analyses beyond body weight focused on this time window. In PPG^Gq^ mice, repeated PPG neuron activation did not significantly suppress 4h food intake beyond day 4, with the anorexigenic effect progressively deminishing after day 1 (Fig 4e). Consequently, cumulative food intake over the 14-day period did not differ significantly between groups (Fig 4d). In line with this, body weight in PPG^Gq^ mice was reduced on day 1 (Fig 4d), but the difference from PPG^Control^ mice was not statistically significant and diminished over time (Fig 4d). A similar transient effect was observed for water intake and EE (Fig 4f,g), with no significant group differences detected after day 1. In contrast, respiratory exchange ratio (RER) remained significantly lower in the PPG^Gq^ group in response to each CNO injection throughout the treatment period, confirming that daily chemogenetic activation has sustained effects which are sufficient to promote higher fat utilisation (Fig 4h). Together, these findings suggest that in the non-obese state, sustained PPG neuron activation only has transient effects on food intake and body weight regulation.

### Chronic PPG neuron activation sustainably reduces food intake and body weight in obese mice

To assess the long-term efficacy and metabolic benefits of brainstem PPG neuron activation in the context of obesity, we repeated the chronic activation experiment in a more translationally-relevant model of diet-induced obesity. PPG^Gq^ and PPG^Control^ mice were fed a 45% high-fat diet *ad libitum*, resulting in a substantial increase in body weight (PPG^Gq^: 21.19+/-2.77%; PPG^control^: 26.28+/-3.39%) and significantly impaired glucose tolerance in both groups after six weeks (Suppl Fig 5a,b). No significant virus x diet interaction was detected in metabolic parameters (Suppl Fig 5c,d), indicating that metabolic changes were driven solely by the high-fat diet, without any confounding constitutive effect of hM3Dq expression prior to the commencement of chronic CNO administration. Obese mice consumed more food and less water during the dark phase, whilst their EE was increased and RER decreased in both light and dark phases (Suppl Fig 5c,d). During the dark phase, satiety was not impaired as meal number and inter-meal interval remained unchanged. However, the overall increased caloric intake in obese mice was driven by calorifically larger meals (Suppl Fig 5c), despite their shorter duration, explained by the higher energy density of high-fat diet. This indicates that impaired satiation is the primary driver of weight gain in this common model of diet-induced obesity, suggesting that activating PPG neurons may represent a rational strategy to counter this driver of overconsumption.

We thus tested the effects of chronic PPG neuron activation in mice with an established diet-induced obese state. Mirroring effects in the lean state, PPG^Gq^ mice showed a marked reduction in caloric intake on the first treatment day (Fig 5a), driven by increased satiation and satiety (Fig 5b), along with decreased water intake and RER (Suppl Fig 4b). However, in the obese state, food intake remained significantly lower for a longer duration (up to hour 7, Suppl Fig 4b), and only modest reductions in EE were observed. Consequently, a significant reduction in body weight was detected 24 hours after CNO injection (Fig 5c).

**Figure 5.**
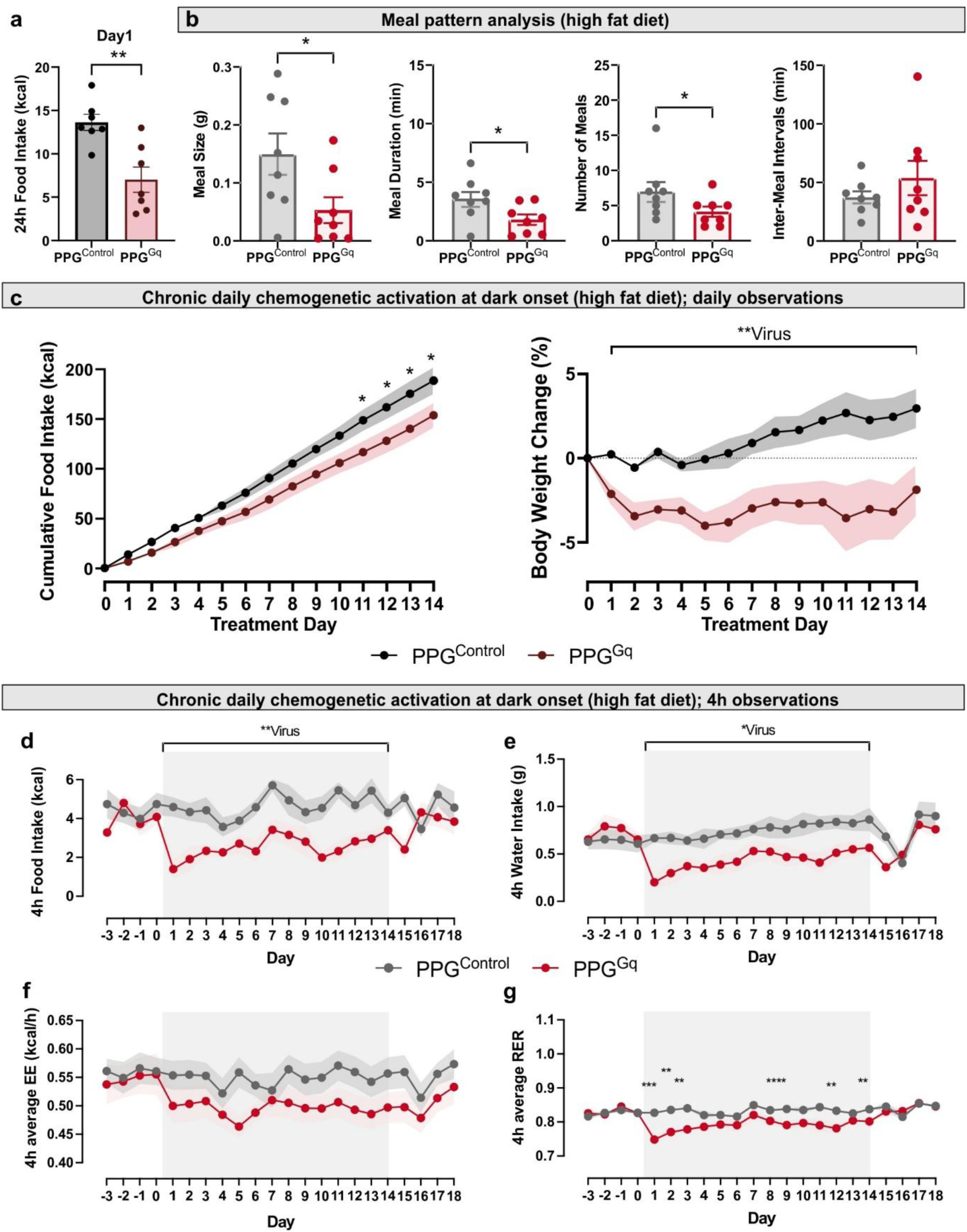
In the obese state repeated chemogenetic activation of PPG^NTS^ and PPG^IRT^ neurons significantly decreased food intake, water intake and body weight in a sustained manner. PPG^Gq^ (n=8) and PPG^Control^ (n=8) mice maintained on high-fat diet for six weeks and then introduced into the Promethion Core^TM^ system. **a** 24h cumulative food intake after the first CNO injection was significantly reduced in PPG^Gq^ compared to PPG^Control^ mice. **b** Meal pattern analysis demonstrating significant increase over the dark phase revealed significant changes in satiation and satiety parameters. **c** Cumulative food intake and body weight change over 14 days of repeated daily CNO (2mg/kg, i.p.) administration 30 min before dark onset in PPG^Gq^ and PPG^control^ mice. 2-way ANOVA, Time x Virus interaction: F(14,168)=3.62, ****p<0.0001. Body weight change (%) was calculated by normalising to the body weight on the day before treatment started (day-1). 2-way ANOVA, Time x Virus interaction: F(13,167)=1.43, p=0.2; Virus: F (1,14)=11.77, **p=0.004. Effects on food intake (**d**), water intake (**e**), average EE (**f**) and average RER (**g**) over the first 4h after each daily CNO administration. Food intake over 4 hours. 2-way ANOVA, Time x Virus interaction: F(13,166)=1.18, p=0.3, Virus: F(1,14)=25.03, ***p=0.0002. Water intake over 4 hours. 2-way ANOVA, Time x Virus interaction: F(13, 167)=0.57, p=0.9, Virus: F(1,14)=6.86, *p=0.02. Average respiratory exchange ratio (RER): 2-way ANOVA: Time x Virus: F(13,166)=2.57, **p=0.003. Fisher’s LSD test: Day1: ***p=0.0002, Day2: **p=0.002, Day3: ****p<0.0001, Day9: **p=0.005, Day12: **p=0.005, Day14: **p=0.007. Average hourly energy expenditure. Data analysed using ANCOVA adjusted for body weight for each treatment day with a significance threshold set at p value<0.01. 2-way ANOVA was conducted for the 14 treatment days, followed by Fisher’s LSD post-hoc test with a significance threshold set at p value<0.01. All data are presented as mean±SEM.

**Figure 6.**
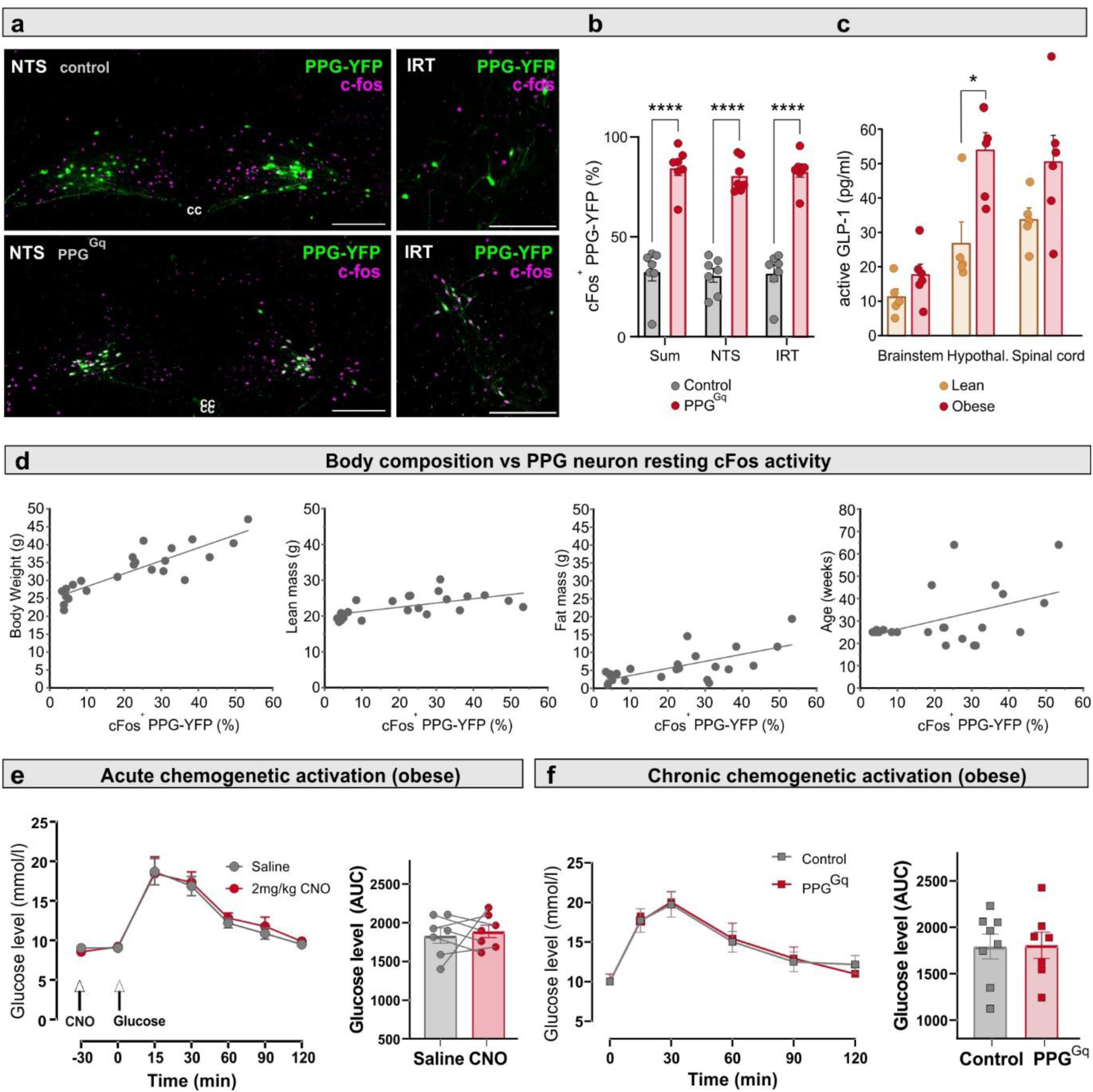
Body composition correlates with PPG neuron activity and obese animals store more GLP-1 within the brain. a Photomicrographs showing expression of cFos immunofluorescence and YFP-expressing PPG neurons in NTS and IRT of PPG^Control^ (top) and PPG^Gq^ (bottom) mice after CNO administration. Scalebars: 200μm. cc, central canal. **b** Quantification of cFos expression in YFP^+^ PPG neurons in PPG^Gq^ and PPG^Control^ mice. **c** Active GLP-1 tissue content in brainstem, hypothalamus and spinal cord of lean (chow diet; n=5) or obese (high-fat diet; n=6) PPG-Cre-YFP mice. *p=0.017. **d** Body composition (body weight, lean mass, fat mass) and age of 23 randomly selected PPG-Cre-YFP mice plotted against the percentage of cFos^+^ PPG neurons in NTS and IRT. Linear fits were produced. The percentage of cFos^+^ PPG neurons was positively correlated with body weight (slope = 0.38 ± 0.10, p = 0.001, R^2^ = 0.41), lean mass (slope = 0.12 ± 0.037, p = 0.004, R^2^ = 0.35) and fat mass (slope = 0.20 ± 0.049, p = 0.0005, R^2^ = 0.46). **e,f** Neither acute (**e**) nor chronic (**f**) chemogenetic activation of brainstem PPG neurons affected glucose tolerance as assed by glucose tolerance test. For all the intraperitoneal (i.p.) glucose tolerance tests (IPGTT), mice were fasted for 5 hours prior to the test. Glucose (1g/kg; i.p.) was given at the start of the test (time 0) and the glucose level was measured from the tail vein blood immediately after glucose administration and at 15, 30, 60, 90 and 120 min after. The acute test was performed with a within subjects design on PPG^Gq^ mice (n=7), whilst the chronic test was between subjects, comparing PPG^Control^ (n=8) with PPG^Gq^ (n=7) mice. In both tests area under the curve was calculated and was found not significantly different between groups. Data presented as mean±SEM.

Chronic PPG neuron activation over 14 days in obese PPG^Gq^ mice led to a significant and sustained reduction in food intake and body weight compared to obese PPG^Control^ mice (Fig. 5c). The hypophagic effect was reflected in cumulative intake, with PPG^Gq^ mice exhibiting significantly lower cumulative intake than controls from day 11 (Fig. 5c). PPG^Gq^ mice also showed persistently reduced 4h water intake throughout the treatment period, likely reflecting decreased eating-associated drinking (Fig 5d). As in the lean state, 4h RER in PPG^Gq^ mice was most strongly decreased on the first day and remained significantly lower than controls for most of the treatment period. While 4h EE appeared lower in PPG^Gq^ mice, the group differences were not statistically significant. These data demonstrate that PPG neurons in the obese state retain their responsiveness and that the resulting hypophagia and weight loss is not compromised but enhanced compared to the lean state.

### Intrinsic PPG neuron activation positively correlates with body weight and fat mass

PPG^Gq^ and PPG^control^ mice on HFD underwent a final CNO administration followed by cFos immunolabelling to assess neuronal activation under HFD to validate that PPG neurons retain responsiveness to chemogenetic activation in the obese state and after the chronic stimulus paradigm. PPG^Gq^ mice had significantly higher levels of cFos-expressing PPG neurons following CNO administration compared to PPG^control^ mice (Fig 6 a,b). Interestingly, we noticed a distinctly higher percentage of cFos-expressing PPG neurons in PPG^control^ mice compared to our previous findings in lean mice^12, 27, 28^. We hypothesised that this increased intrinsic activity of PPG neurons may reflect an obesity-induced upregulation of central GLP-1 signalling which we tested by analysing GLP-1 content across different brain regions in mice on standard chow or HFD. This revealed significantly elevated GLP-1 content in the hypothalamus (a major PPG neuron projection target) in the obese state, with a trend for increase also observed in brainstem and spinal cord (Fig 6c).

To further explore whether PPG neuron activity correlates with body composition parameters, we determined the percentage of cFos^+^ PPG neurons in a cohort of naive mice varying in body composition, sex and age (Fig. 6d). PPG neuron activation positively correlated with body weight, lean mass and fat mass. This correlation was stronger for fat (Pearson’s r: 0.68) than for lean mass (Pearson’s r: 0.59). Although age also showed a positive correlation with PPG neuron activation (Fig. 6d), this effect was largely accounted for by its covariance with fat mass (Suppl Fig 6). Altogether, this indicates that, adiposity rather than lean mass, might be the main driver for PPG neuron activity, possibly via leptin signalling^15^. Alternatively, body weight itself (Pearson’s R: 0.87), as proposed by the gravitostat theory^29^, might be the driver, although the mechanism by which it would activate PPG neurons is unclear.

### Neither acute nor chronic activation of PPG neurons improves glucose tolerance in the obese state

While gut-secreted GLP-1 is widely known to improve glucose tolerance, the role of brain-derived GLP-1 in peripheral glucose regulation remains a subject of controversy. PPG^NTS^ neuron activation has been reported to either improve glucose tolerance^30^ or show no effect^14^ and we previously failed to see an effect on glucose tolerance when PPG^NTS^ neurons were ablated^13^. Here, we focused on the obese state when impaired glucose tolerance has already developed (suppl Fig 5a,b) and assessed whether combined activation of both PPG^NTS^ and PPG^IRT^ neurons alters glucose tolerance. Firstly, we investigated the effect of acute PPG neuron activation in an intraperitoneal glucose tolerance test (IPGTT). PPG neuron activation had no impact on fasting glucose levels 30 minutes after CNO injection, and there was no change in the glucose area under the curve (AUC, Fig 6e). Next, we asked whether chronic PPG neuron activation improves glucose tolerance. To do this, an IPGTT test was performed one day after 14-day repeated PPG neuron activation, in the absence of acute PPG neuron activation. There was no significant difference in either fasting glucose levels or glucose tolerance between PPG^Gq^ and PPG^Control^ mice (Fig. 6f). These findings suggests that neither acute nor chronic PPG neuron activation influences peripheral glucose homeostasis.

Altogether, our studies demonstrate that PPG^NTS^ and PPG^IRT^ neurons integrate distinct inputs in a convergent manner to modulate physiological satiation. Crucially, these neurons have the capacity to potentiate both satiation and satiety when supraphysiologically activated, leading to sustained weight loss in the obese state, providing a strong rationale for this population as a target for obesity pharmacotherapy.

## Discussion

This study produced important new insights into the physiological importance and utility for obesity treatment of brainstem PPG neurons. Firstly, we examined the functional anatomy of brainstem PPG neurons. By comparing the connectivity of PPG^NTS^ and PPG^IRT^ neurons, we demonstrated they receive largely distinct inputs, most notably an absence of vagal afferent inputs to PPG^IRT^ neurons but have very similar projection patterns. In addition, we showed an absence of crosstalk between these populations of PPG neurons using both optogenetic and chemogenetic approaches. This circuit organisation suggests that PPG neurons from both populations serve to integrate diverse input signals and convey these to multiple common postsynaptic targets in a largely parallel convergent and divergent circuit structure. Consequently, the influence of PPG neurons on food intake is a delicate balance between a large array of parameters being considered. Each individual PPG neuron contributes its activity based on its specific sets of inputs to this balance. Experimentally, only when the balance between active and non-active PPG neurons is strongly tipped in one direction, do we detect instantly measurable effects food intake. Such a situation arises when either virtually all PPG neurons are activated (e.g. chemogenetically) and thereby produce a highly significant reduction in food intake, or when almost all PPG neurons are ablated as in this study, which then leads to overeating and slow excessive weight gain. This relatively small contribution of each cell only becomes apparent as body weight gain or loss over longer timescales and in the various correlations between number of functional or active PPG neurons and body parameters that we observed across a population of mice, such as the fact that the number of active PPG neurons is an indicator of body mass.

The second major finding from the current study is that persistent weight loss and hypophagia can be achieved by chronic PPG neuron activation in the obese but not the lean state. This went hand in hand with the observation that baseline PPG neuron activity is increased and that the amount of GLP-1 detected in brain tissue is increased under HFD. Similarly, in rat a higher level of brainstem PPG mRNA expression was found in rats with greater fat mass^31^ and in obese Zucker rats^32^. We also demonstrated recently the feasibility of pharmacological approaches to activate PPG neurons with the 5-HT₂C receptor agonist lorcaserin^10^. These observations demonstrate that indeed pharmacological activation of the central GLP-1 system is a promising avenue for enhanced anti-obesity treatment, because the system remains functional in the obese state, in fact, with increased body mass or fat mass, more PPG neurons become active.

The results from the current study demonstrate that PPG^NTS^ and PPG^IRT^ neurons are a driver of *ad libitum* satiation and are also potential targets for anti-obesity therapies. PPG neuron activation reduced body weight mainly through increasing satiation and satiety, and it also shifted nutrient partitioning towards greater fat utilisation. We show that ’physiological’ activation by diet-induced obesity only activates less than half of all PPG neurons and thereby only slightly ameliorates weight gain, as also demonstrated by the effect of PPG neuron ablation, but that full activation of this cell population, here achieved by chemogenetic activation, has the capacity to substantially reduce food intake. This suggests that PPG neurons are strategically positioned to be influenced during obesity development, making them a promising anti-obesity drug target. In the current study, a tendency of reduction in energy expenditure was observed, which potentially reduced the net anti-obesity effect of PPG neuron activation. This could be improved by combining PPG neuron activation with drugs which augment energy expenditure such as glucagon receptor agonists^33^. Additional research is warranted to explore further approaches and drugs for PPG neuron activation, and to investigate potential combination therapies, with a specific emphasis on modalities that augment energy expenditure.

## Methods

### Animals

Adult transgenic C57BL/6NJ mice (8-34 weeks old) of both sexes expressing Cre recombinase or yellow fluorescent protein (YFP) under the glucagon promotor (PPG-Cre or PPG-YFP^34, 35^) were used in this study. These two strains were also crossed to obtain PPG-Cre-YFP mice. Mice were fed with either normal chow (Teklad Global Rodent Diets, 2018C, 3.1kcal/g) or high-fat diet (D12451i; Research Diets, USA, 4.73kcal/g) containing 45% kcal fat and were provided with water *ad libitum*. Mice were kept under 12-hour light/dark cycle (light off at 2pm) at 20-24℃ and humidity of 45-65%. All procedures were performed in accordance with the UK Animals (Scientific Procedures) Act 1986 and with approval from the UCL Animal Welfare and Ethical Review Body.

### Stereotaxic Surgery and viral gene transfer

Stereotaxic surgery was performed as described previously^12, 13^. Briefly, mice were anesthetised using ketamine hydrochloride (Ketavet®, Zoetis; 75 mg/kg, i.p.) and medetomidine hydrochloride (Domitor®, OrionPharma; 1 mg/kg, i.p.), with meloxicam (Metacam®, Boehringer Ingelheim; 5mg/kg, s.c.) for peri-surgery analgesia. Mice were mounted onto a stereotaxic frame, with their head bent downwards to be perpendicular to the bench surface. An incision was made in the scalp to expose the atlanto-occipital membrane covering the area between the occipital crest and the first vertebra. The meninges were cut open using a 30G needle to expose the brainstem surface. A volume of 200-250nl virus (detailed in Table 1) per injection site was delivered at 50nl/min using a pressurised glass capillary. Coordinates were (in reference to obex): 0.1 mm rostral, ±0.5 mm lateral, -0.35 mm ventral for NTS targeting; and -0.5 mm rostral, ±1.1 mm lateral, -1.4mm ventral for IRT targeting. Body temperature was constantly monitored using an animal temperature controller (WPI, ATC2000). Atipamezole hydrochloride (Antisedan®, OrionPharma; 1mg/kg, s.c.) was given for anaesthesia reversal. Mice were allowed at least 3 weeks recovery before start of experiments.

**Table 1.**
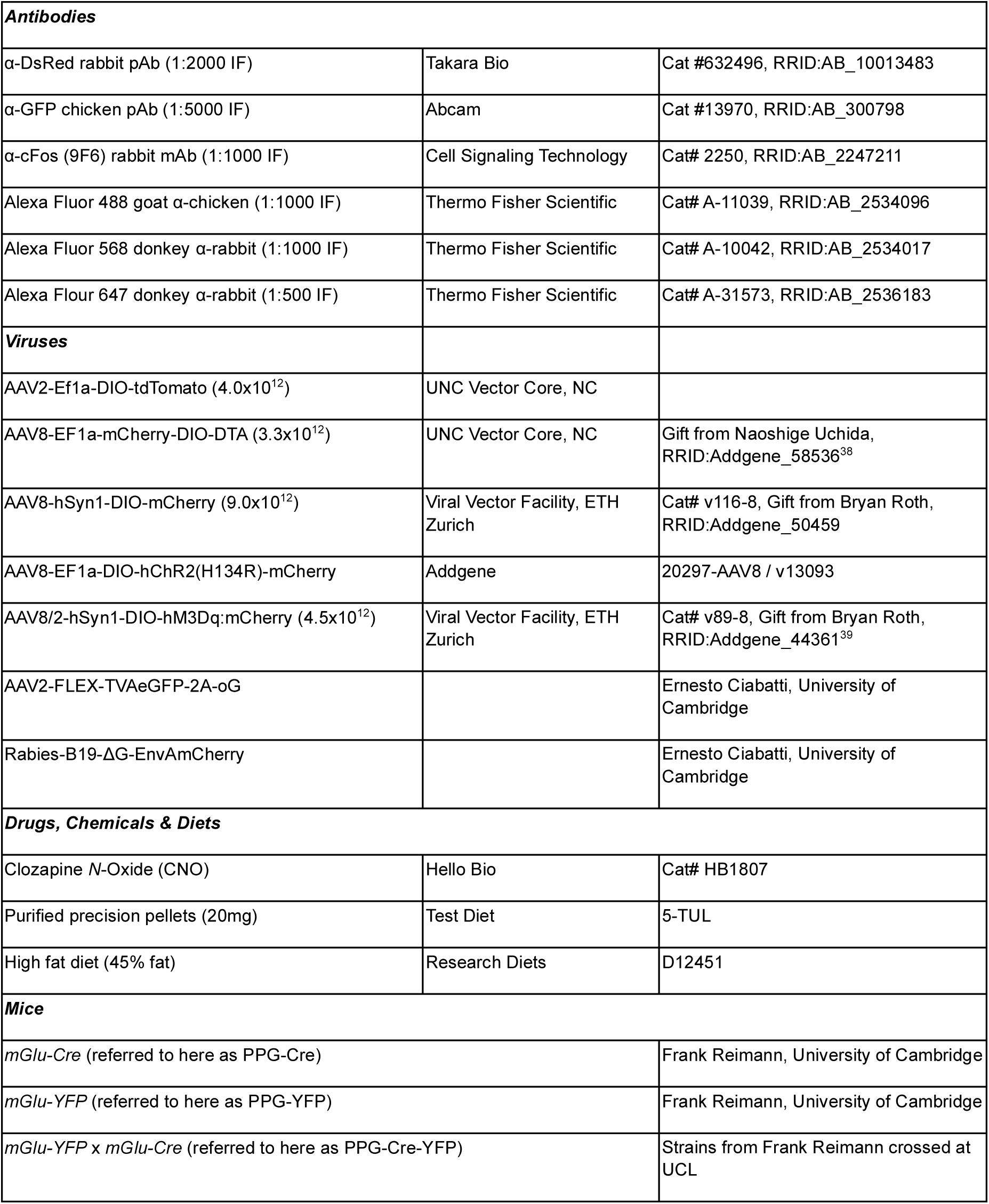
Reagents and Resources.

### Monosynaptic retrograde tracing of PPG neurons

Monosynaptic retrograde tracing was performed as previously described in the study of Lewis et al. (2022). To assess if PPG^NTS^ and PPG^IRT^ neurons receive monosynaptic inputs from the same anatomical locations, PPG-Cre-YFP mice (n=5) were injected bilaterally into either the NTS or the IRT with 200 nl of a Cre-dependent tri-cistronic helper virus (AAV2-FLEX-TVAeGFP-2A-oG), enabling Cre-dependent expression of the TVA receptor, green fluorescent protein (GFP), and rabies glycoprotein (oG) specifically in PPG neurons. 21 days post injection, mice received a unilateral injection of 500 nl of EnvA-pseudotyped, glycoprotein-deleted rabies virus encoding mCherry (Rabies-B19-ΔG-EnvAmCherry) into the same region. The TVA receptor in PPG neurons facilitates infection by the EnvA-pseudotyped, G-deleted rabies virus, while the coexpressed oG enables retrograde transsynaptic spread to monosynaptically connected neurons. This approach labels both PPG neurons and their monosynaptic input neurons with mCherry fluorescence. Seven days after rabies virus injection, mice were transcardially perfused for tissue harvesting and IHC processing. Additionally, a PPG-YFP mouse lacking Cre recombinase expression (PPG-Cre ^-/-^) underwent the same procedure as a negative control.

### Optogenetic activation of PPG^NTS^ neurons

For optogenetic activation of PPG^NTS^ neurons, mice transduced bilaterally with AAV8-EF1a-DIO-hChR2(H134R)-mCherry-WPRE-HGHpA in the NTS only were fasted for 2 hours and then anesthetized using 1.5-2.5% isoflurane, followed by surgical exposure of the NTS and optical stimulation using a laser-coupled optical fibre with 450nm light pulses for 5 min (50ms light at 10Hz, 20mW). 90 min after stimulation mice were deeply anesthetized using isoflurane overdose, followed by transcardial perfusion and tissue processing for c-fos and fluorescent reporter immunofluorescence.

### Chemogenetic activation of PPG neurons

Adult PPG-Cre mice of either sex received bilateral viral injection of AAV8-hSyn1-DIO-hM3Dq:mCherry (PPG^Gq^, n=8) for chemogenetic activation or AAV2-Ef1a-DIO-tdTomato as controls (PPG^Control^, n=8) in both the NTS and IRT regions (Fig 3a,b). Expression of hM3Dq did not affect food intake, water intake, energy expenditure, RER or ambulatory movement in light or dark phase (Suppl Fig 5c,d), nor did it affect meal patterns (Suppl Fig 5c). To assess the effect of chronic chemogenetic activation of PPG neurons a dose of 2 mg/kg CNO was injected i.p. once daily 30min prior to dark onset.

For unilateral chemogenetic activation of NTS PPG neurons, adult PPG-Cre-YFP mice of either sex received unilateral NTS injection of AAV8-hSyn1-DIO-hM3Dq:mCherry (PPG^Gq^, n=5) for chemogenetic activation or AAV2-Ef1a-DIO-tdTomato as controls (PPG^Control^, n=6).

### Immunohistochemistry (IHC)

For validation of PPG neuron ablation using AAV8-EF1a-mCherry-DIO-DTA or transduction of AAV2-Ef1a-DIO-tdTomato, mice were transcardially perfused and the tissue was processed with IHC for amplification of fluorescent reporter signals. To validate chemogenetic activation and viral transduction of AAV8- hSyn1-DIO-hM3Dq:mCherry and AAV2-Ef1a-DIO-tdTomato virus, mice were administered with either saline or 2mg/kg CNO (i.p.). 90 min later mice were deeply anesthetised using pentobarbital (Vetoquinol, 100 mg/kg) and perfused transcardially with ice-cold 0.1M Phosphate Buffer (PB), followed by 4% paraformaldehyde (PFA) in 0.1M PB. Brains were extracted and post-fixed in 4% PFA overnight, followed by cryo-protection using 30% sucrose in 0.1M PB. 30 µm thick coronal brain sections were cut using a cryostat, and stored in cryoprotectant (30% sucrose, 1% Polyvinyl-pyrrolidone and 30% ethylene glycol in 0.1M PB) until further use.

IHC was performed using free-floating sections as previously described^13^. For amplification of fluorescent reporter YFP, tdTomato and mCherry, sections were first incubated with 0.5% sodium borohydride (Sigma Aldrich 71320) in 0.1 M PB for 20min to reduce autofluorescence. Sections were then incubated with rabbit anti-DsRed (1:2000) and chicken anti-GFP (1:5000) primary antibodies overnight in blocking solution (0.1M PB containing 0.3% Triton, 0.5% normal goat serum, 0.5% donkey serum), followed by incubation with Alexa Fluor 488 goat α-chicken (1:1000) and Alexa Fluor 568 donkey α-rabbit (1:1000) secondary antibodies in blocking solution for 2 hours in the dark. Sections were washed for three times before and after the IHC processing, and in between each IHC step.

For c-fos IHC, free-floating sections were treated with 10 mM sodium citrate (pH 6.0; Alfa Aesar, A12274) at 80 °C for antigen retrieval, followed by washes with 0.1M PB. Next, sections were incubated with blocking solution (0.1M PB containing 5% donkey serum and 0.3% Tween20 (Millipore 655204-100mL)) for 1 hour, followed by overnight incubation with rabbit anti-c-fos primary antibody (1:1000) in blocking solution. Following this, the sections underwent three 10-min washes with 0.1% Tween20 in 0.1M PB, followed by 2-hour incubation in the dark with AlexaFluor647 donkey anti-rabbit (1:1000) secondary antibody prepared in blocking solution. All antibodies used are detailed in Table 1. After IHC processing, brain sections were mounted on slides, coverslipped with VECTASHIELD Antifade MountingMedium (H-1200) and imaged using an EVOS M7000 microscope (Thermo Fisher Scientific) or a Leica SP8 confocal microscope. Images were edited using Fiji software^36^.

### Food Intake Studies

FED3 pellet dispensers^37^ were used for experiments assessing the effect of ablation or acute activation of PPG neurons whilst a Promethion Core^TM^ (Sable Systems) metabolic and behavioural phenotyping system was used to monitor the effects of repetitive activation of PPG neurons. Studies were conducted using a between-subject design with sex and age balanced groups unless otherwise stated.

FED3 pellet dispensers were fitted into home cages and normal chow 20mg Precision Pellets (TestDiet, USA, 5TUL) were provided as food source. Mice were fed *ad libitum* with a single pellet dispensed within 5 seconds after the previous pellet was taken from the pellet well. Mice were habituated to the FED3s until stable food intake was reached for at least 3 consecutive days. To assess the effect of PPG neuron ablation on food intake under baseline conditions, mice were left undisturbed for 3 sessions of 48 hours, with a one-day interval between sessions. On the intervening day, mice were weighed and FED3s were removed for data collection at 11am and returned upon dark onset at 2pm. For assessment of the effect of acute activation of PPG neurons, mice were first habituated to the FED3s and saline injection (i.p.). Mice were considered habituated once food intake was not significantly different across three consecutive sessions of saline injection (i.p.) separated by one wash-out day. Treatment days were separated by two days of washout, during which no injection was given. On the treatment day, mice were fasted for 3 hours before dark-onset and CNO was administered 30 min before dark-onset. Body weight was measured, and food intake data were collected on the following day during the 3h fast.

The Promethion Core system provided continuous measurement of food and water intake, body weight, energy expenditure, respiratory exchange ratio (RER) and locomotor activity. Mice were first habituated to the Promethion Core system and saline injection. Following this, to assess the effect of chronic activation of PPG neurons, mice were left undisturbed for four days. This was followed by daily i.p. injection of CNO (2mg/kg) for 14 consecutive days. Mice were fasted daily for 3 hours before dark-onset and 2mg/kg CNO was administered 30 min before dark-onset. Under high-fat diet, mice were given 2mg/kg CNO for an additional day (15^th^ day) to ensure collection of 24-hour data across the 14-day period, while also allowing assessment of the effect of chronic PPG neuron activation on glucose tolerance the next day (16^th^ day) without cessation of chronic activation. Recordings finished with an additional three or four days of washout without i.p. injection in the obese and lean state, respectively.

### Meal pattern analysis

The meal definition was established using feeding data from 16 mice (10-15 weeks old) recorded using the FED3 system. Given that meal pattern analysis was restricted to the dark phase in the current study, only food intake data from this phase were used for generating the meal definition.

Briefly, the FED3 system registers the time of each pellet removal, with the intervals between pellet collections (inter-pellet intervals, IPIs) categorized as either within-meal intervals (WMIs) or inter-meal intervals (IMIs; Suppl Fig 7a). The IPIs were then log-normalized, and the resulting density plot revealed a bimodal pattern, corresponding to the distributions of WMI and IMI (Suppl Fig 7b), respectively. The IMI threshold, defined as the minimum IPI separating two consecutive meals, was established based on previous literature (Zorrilla et al., 2005; Rathod and Di Fulvio, 2021). To do this, we tested a range of thresholds, calculated the corresponding meal size, duration and number of meals, followed by analysing the rate change of these curves (Suppl Fig 7c). Furthermore, to address the issue of short IMI thresholds excluding pellets from meals, we assessed the percentage of pellets included in meals across various thresholds, with the aim to maximise pellet inclusion (Suppl Fig 7c). As a result, the value of 10^2.24^ sec (about 3min) was chosen as the threshold. This value falls within the range of tested threshold values which gave minimal rate of change in meal size, duration and number of meals, while allowing the maximal possible number of pellets (85%) to be included in a defined meal. Additionally, the same analysis was conducted using feeding data from the Promethion Core system (Suppl Fig 7d) to cross-verify the chosen threshold for meal definition. Finally, due to the technical limitations of the FEDs, which register only the intervals between pellets but not the actual time of pellet consumption, we defined the minimum meal size to be two pellets (0.04g) to allow for the calculation of meal duration. As a result, pellets consumed within 3 minutes of each other were defined as belonging to the same meal, with a minimum meal size set at 0.04 g.

The WMI and IMI distributions were visually identified as resembling lognormal and normal Gaussian distributions. Nonlinear regression analysis was performed in GraphPad prism, followed by Akaike’s information criterion (AIC) analysis to determine the best model of fit. This allowed for the generation of mathematical formula for the IPI density distribution registered by the FED system as follows:

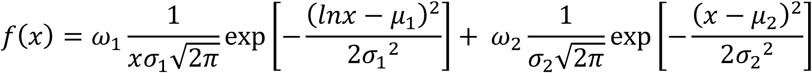

where x is the duration of IPI, with *ω*_1_ = 0.69, *μ*_1_ = 0.13, *σ*_1_ = 0.12; *ω*_2_ = 0.23, *μ*_2_ = 2.93, *σ*_2_ = 0.36.

### EchoMRI

PPG-Cre-YFP mice (n=23) of both sexes (19-64 weeks old) were maintained on an *ad libitum* chow diet. To assess obesity levels, these mice first received a body composition scan (EchoMRI^TM^-100H) to measure lean and fat mass. Immediately after the scan, the mice were perfuse-fixed for brain tissue collection. This procedure was conducted during the late light phase (between 11:00 AM and 2:00 PM) to minimise potential food-related activation of PPG neurons.

### Intraperitoneal glucose tolerance test (IPGTT)

All IPGTTs were conducted during the 2 hours before dark onset. Mice were transferred to new cages and fasted for 5 hours with *ad libitum* access to water prior to the start of the IPGTT. Following this, mice received 1g/kg glucose (20% glucose in sterile saline, i.p.). Blood samples were collected from the tail vein immediately before the glucose injection for baseline glucose measurement (Time 0), and then at 15, 30, 60, 90 and 120 min after the injection. Plasma glucose levels were measured Immediately after collection using a Roche ACCU-CHECK (Aviva) test kit. For the assessment of the effect of chronic PPG neuron activation on glucose tolerance, a between-subject design was used with no drug interventions. When assessing the effect of acute activation of PPG neurons on glucose tolerance, a within-subject design was used, and it consists of two experimental days separated by a 3-day washout period. On each experimental day, 30min prior to the baseline glucose measurement (Time -30), the blood glucose levels were also measured, which was followed immediately by either 2mg/kg CNO (5ml/kg, i.p.) or saline (5ml/kg) administration.

### Active GLP-1 assay

Mice were anaesthetised with isoflurane, brains and spinal cords were rapidly extracted, and brainstem, hypothalamus and thoracic spinal cord microdissected and snap-frozen on dry ice. Subsequently, samples were homogenised in PBS with DPP-4 inhibitor (Millipore) and refrozen. Upon thawing samples were centrifuged twice (500g for 10min) and the supernatant was collected and active GLP-1 content determined with a MSD kit (K150JWC-1; Meso Scale Diagnostics).

### Data analysis and Statistics

For quantification of cell number in the PPG neuron ablation experiment, all sections along the anterior-posterior axis of the brainstem (bregma -8.24 to -7.64) were used for manual counting. In the remaining experiments, cells were counted from one of the four series of sections, which were situated along the same rostro-caudal levels and about 0.12mm apart. Food intake and metabolic data collected from the Promethion Core System were first processed using Macro Interpreter software packages (Sable Systems), followed by processing in R Studio (Version 2022.12.0+353). Data collected from FED3s underwent initial processing in R Studio. All statistical analysis and graph plotting was performed using GraphPad Prism (Version 10.2.0 (335)). All data are presented as mean±SEM. Depending on the data, linear regression analysis, student t-test, nested t-test, one-way ANOVA followed by Bonferroni correction, or 2-way ANOVA followed by Sidak post-hoc analysis or Fisher’s LSD test were used, as appropriate. Statistical results are detailed in the figure legends.

## Acknowledgements

This study was supported by the EFSD European Research Programme on New Targets for Type 2 Diabetes supported by an educational research grant from MSD to ST and DB, a MRC project grant (MR/X003604/1) to ST and FR, and an ALERT21 equipment grant from the BBSRC to ST. MKH was funded by a BHF fellowship and WJ was supported by a UCL ORS scholarship and a CSC scholarship from the Chinese government. The authors would also like to thank Dr Diana Williams and Dr Alan Proctor for valuable discussions on meal pattern analysis.

## Author contributions

The concept was developed by WJ, MKH and ST. The methodology was designed by WJ, CS, DIB, MKH and ST. The experimental work was carried out by WJ, CS, IR and MKH. Resources were provided by FMG, FR, EC, DIB, MKH and ST. Data were curated by WJ, CS, MKH and ST. The original draft was written by WJ and ST. Review and editing of the manuscript were conducted by all authors. Visualization was done by WJ, CS and ST. The work was supervised by ST. Project administration was done by WJ, CS, MKH and ST. Funding was acquired by DIB, MKH and ST. (contributions are defined according to the CRediT taxonomy).

