## Supplemental Figures for "Brainstem GLP-1 neurons modulate physiological satiation and drive sustained weight loss in obese mice"

### Supplemental data

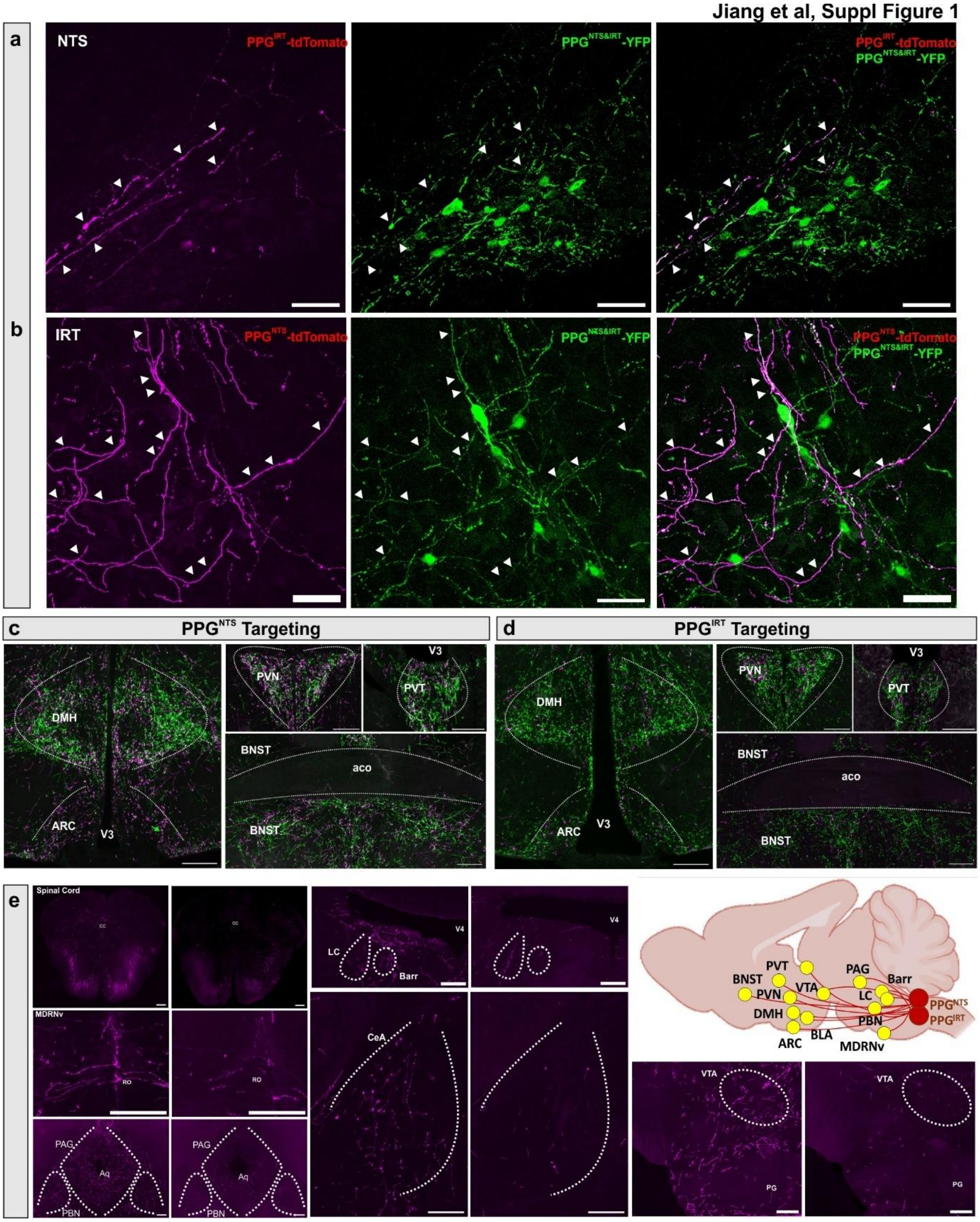


**Suppl Figure 1.** PPG$^{NTS}$ neurons project significantly into the IRT region and PPG$^{IRT}$ neurons project to the NTS. Mice received bilateral injection of AAV2-Ef1a-DIO-tdTomato virus into either NTS or IRT regions to induce tdTomato fluorescence expression in the PPG$^{NTS}$ (n=3) or PPG$^{IRT}$ (n=3) neurons in PPG-Cre-YFP mice. **a,b** Representative images showing colocalization between YFP and tdTomato immunofluorescence in axons originated from PPG$^{NTS}$ or PPG$^{IRT}$ neurons. Scale bars: 50 μm. **c,d** Photomicrograph showing tdTomato-immunoreactive axons originated from $PPG^{NTS}$ and $PPG^{IRT}$ neurons, respectively (magenta), compared to all PPG neuron projection (green) in various brain areas. Scale bars: 200µm. **e** Further brain areas receiving projections from $PPG^{NTS}$ (left) and $PPG^{IRT}$ (right) neurons, respectively. Scale bars: 200µm. Schematic shows brain areas receiving projections from both $PPG^{NTS}$ and $PPG^{IRT}$ neurons CC: central canal. MDRNv: medullary reticular nucleus, ventral, RO: nucleus raphe obscurus, LC: locus coeruleus, Barr: Barrington's nucleus, V4: $4^{th}$ ventricle, BLA: basolateral amygdala, PAG: Periaqueductal gray, PBN: Parabrachial nucleus, Aq: cerebral aqueduct, VTA: ventral tegmental area, PG: pontine gray, CeA: central amygdala, NTS: nucleus tractus solitarius, IRT: intermediate reticular nucleus.

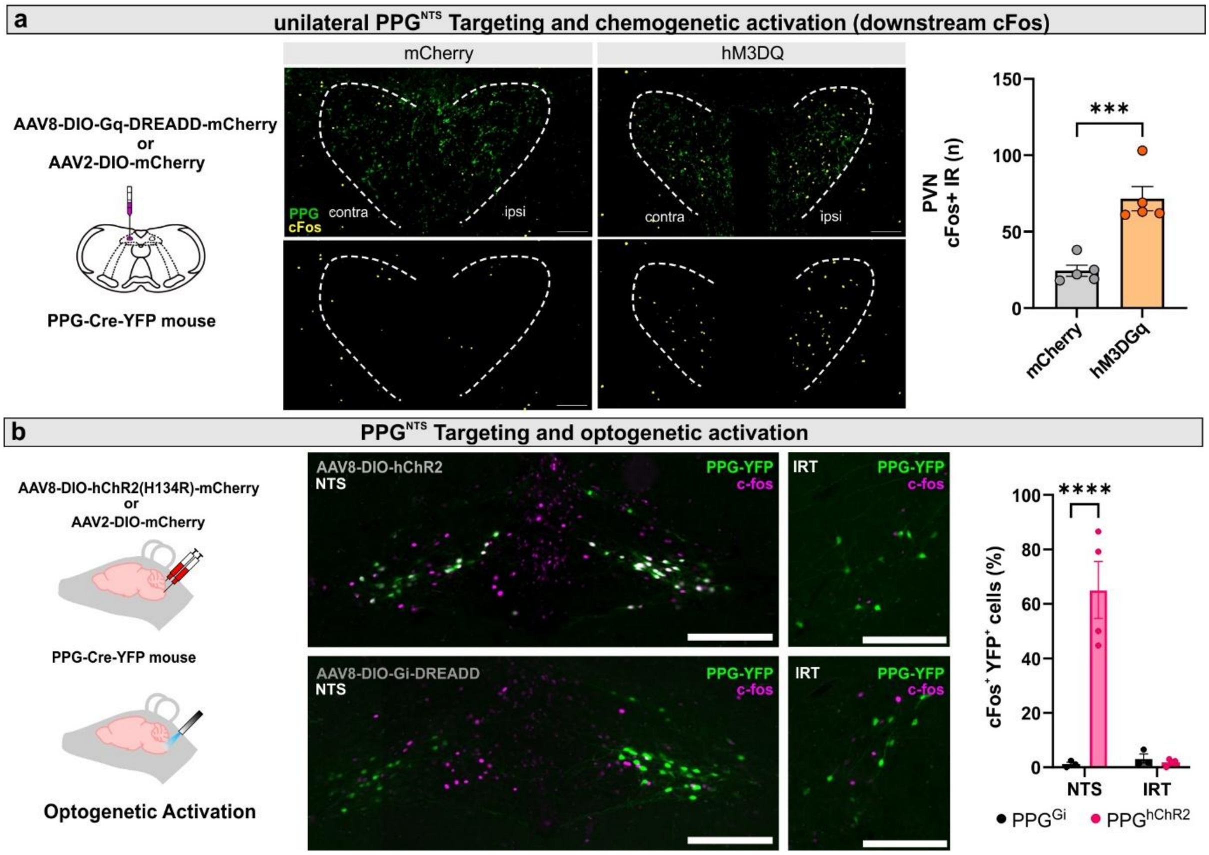


**Suppl Figure 2. a** PPG-Cre-YFP mice received unilateral injection of AAV8/2-DIO-Gq-DREADD or AAV2-DIO-mCherry virus into the NTS (n=5, each) for subsequent unilateral activation via i.p. CNO injection. Middle: representative photomicrographs of cFos expression 90 min after i.p. CNO administration in the PVN of control (mCherry) or $PPG^{Gq}$ mice. Right: quantification of these data. **b** PPG-Cre-YFP mice received bilateral injection of AAV8-DIO-hChR2 or AAV2-DIO-mCherry virus into the NTS (n=4, each) for subsequent unilateral optogenetic activation after fibre implantation. Middle: representative photomicrographs of cFos expression 90 min after unilateral optogenetic stimulation of the NTS of control (mCherry) or $PPG^{ChR2}$ mice. Right: Quantification of cFos-ir positive $PPG^{NTS}$ and $PPG^{IRT}$ neurons and NG. Scale bars: 100 µm.

Jiang et al, Suppl Figure 3

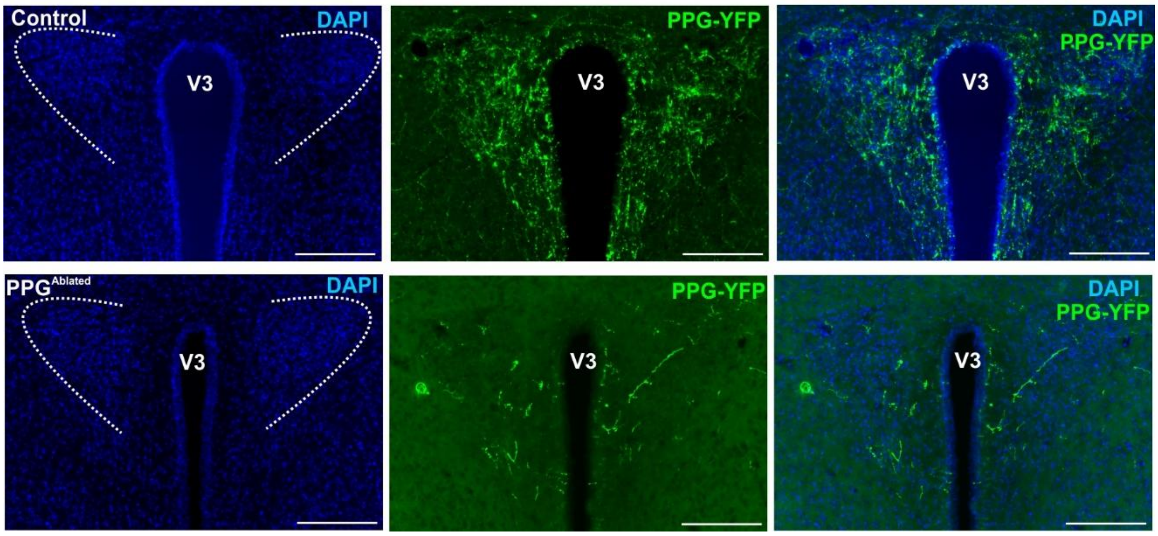


**Suppl figure 3.** Loss of axons in the paraventricular nucleus (PVN) in $PPG^{DTA}$, but not $PPG^{Control}$ mice. PPG-Cre-YFP mice received bilateral viral injections into both NTS and IRT for either PPG neuron ablation ($PPG^{DTA}$, n=8) or tdTomato fluorescence expression as controls ($PPG^{Control}$, n=8). Ablation of PPG neurons in both the NTS and IRT led to almost a complete loss of PPG neuron axons in the PVN. Scale bars: 200µm.

**Jiang et al, Suppl Figure 4**

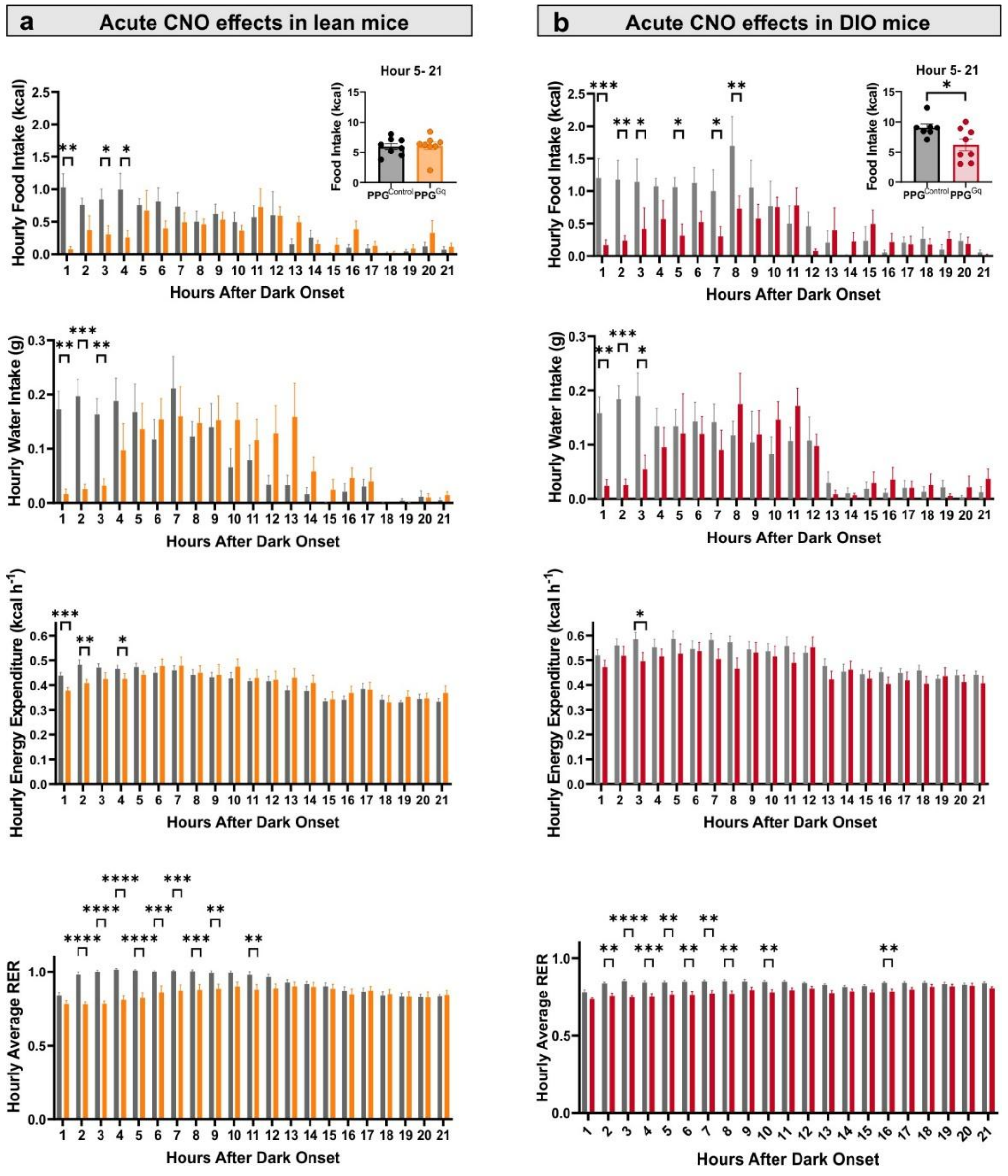


**Suppl figure 4.** The effect of acute activation of $PPG^{NTS}$ and $PPG^{IRT}$ neurons in lean and obese mice. PPG-Cre mice received bilateral injections of either a control virus (AAV2-DIO-tdTomato, $PPG^{Control}$, n=8) or AAV8-DIO-hM3Dq-mCherry ($PPG^{Gq}$, n=8), targeting both the NTS and IRT. Both groups received Clozapine N-oxide (CNO, 2 mg/kg, i.p.) 30 minutes before dark onset on the day of the experiment. Experiments were performed when mice were maintained on a chow diet (**a**) and after six weeks on a 45% high-fat diet (**b**). Food intake, water intake, energy expenditure and respiratory exchange ratio were measured continuously and plotted in hourly bins for the first 21h from dark onset (12h dark, followed by 9 hours of the light phase). ANCOVA was performed on energy expenditure at each time point, and two-way ANOVA was used for all other datasets, followed by Fisher's LSD post-hoc test. The significance level for statistical comparisons conducted at each time point was set at $p < 0.01$. An unpaired t-test was conducted for the total food intake following the first four hours of dark phase. **a** In lean mice, acute activation of PPG neurons significantly reduced hourly food intake (Two-way ANOVA, Time x Virus: $F(23,322)=2.43$, $p=0.0004$) without evidence of compensatory intake later in the day. Significant reduction in hourly water intake. Two-way ANOVA, Time x Virus: $F(20,280)=3.28$, $p<0.0001$. Reduction in hourly energy expenditure. ANCOVA adjusted for body weight. Hour 1: body weight $F(1,13)=11.72$, $p=0.005$; virus $F(1,13)=22.25$, $p=0.0004$. Hour 2: body weight $F(1,13)=5.04$, $p=0.04$; virus $F(1,13)=13.87$, $p=0.003$. Hour 4: body weight $F(1,13)=8.35$, $p=0.01$; virus $F(1,13)=4.78$, $p=0.05$. Significant decrease in hourly respiratory exchange ratio (RER). Two-way ANOVA, $F(20,280)=8.79$, $p<0.0001$. **b** In obese mice, acute activation of PPG neurons significantly reduced hourly food intake in the first 4 hours (Two-way ANOVA, Time x Virus interaction: $F(20,260)=2.21$, $p=0.0025$) and after. Significant reduction in hourly water intake. Two-way ANOVA, Time x Virus interaction: $F(20,260)=2.279$, $p=0.002$. Reduction in hourly energy expenditure. ANCOVA adjusted for body weight, with significance set at $p<0.01$. Body weight: $F(1,12)=14.02$, $p=0.003$; Virus: $F(1,12)=5.04$, $p=0.04$. Significant decrease in respiratory exchange ratio (RER). Two-way ANOVA, Time x Virus interaction: $F(20,260)=3.17$, $p<0.0001$. All data are presented as mean ± SEM. *$p < 0.05$, **$p < 0.01$, ***$p < 0.001$, ****$p < 0.0001$.

Jiang et al, Suppl Figure 5

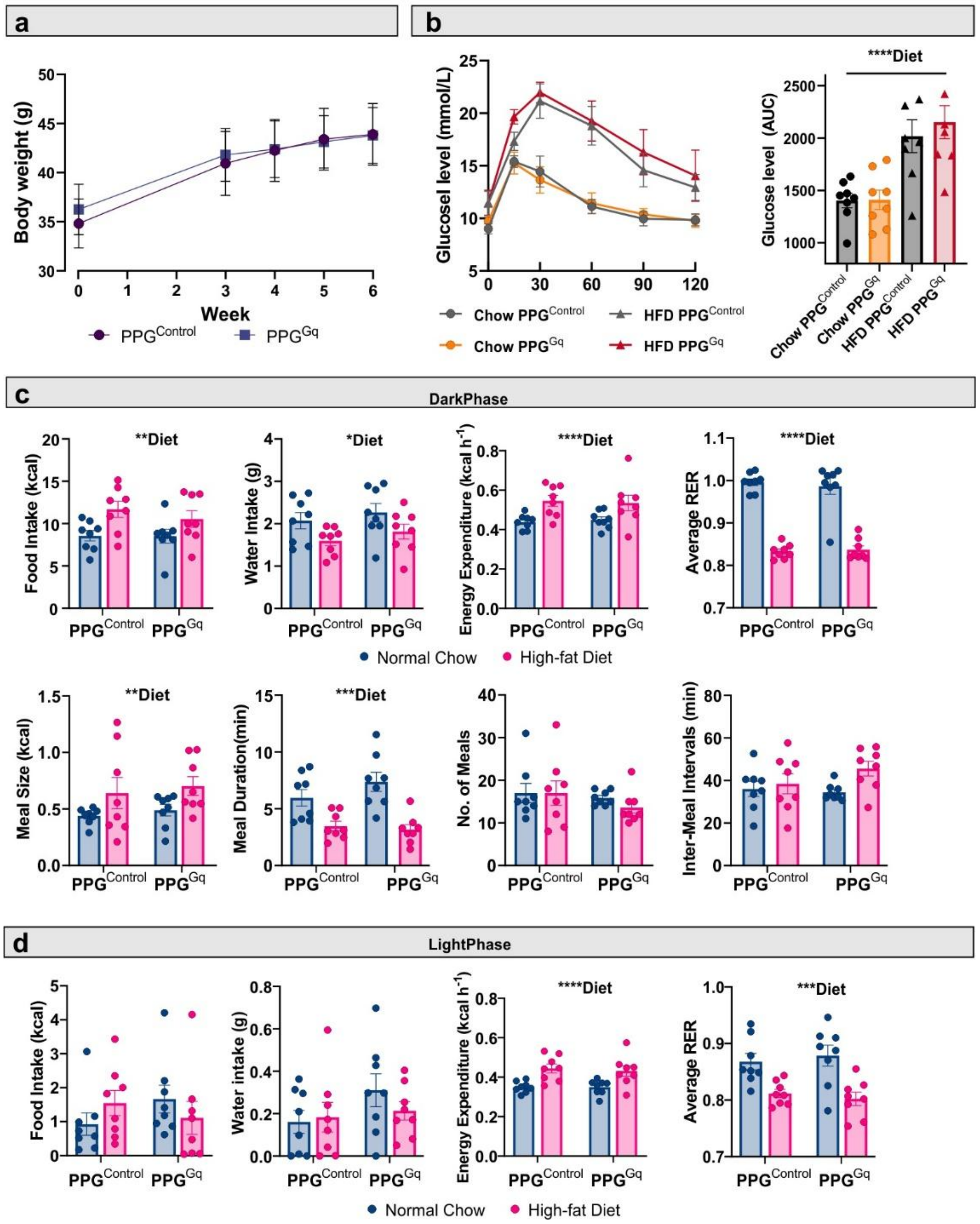


**Suppl figure 5.** PPG$^{Gq}$ and PPG$^{Control}$ mice were fed *ad libitum* a 45% high fat diet (HFD) until a stable body weight was achieved (<5% body weight gain compared to the previous week; **a**). **b** After being fed HFD for 6 weeks, both groups of mice showed glucose intolerance compared to their condition prior to HFD feeding (chow) as determined by i.p. glucose tolerance test. Left: glucose level at various timepoints after glucose (1g/kg; i.p.) injection. Right. Area under the curve (AUC) glucose tolerance for each group). **c** Dark phase food intake, water intake, energy expenditure and average respiratory exchange ratio (RER), for both groups of mice before (Normal Chow) and after (High-fat Diet) 6 weeks of HFD feeding. **d** plots of the same parameters during the light phase. Data obtained after mice were habituated to the Promethion Core™ (Sable Systems) metabolic and behavioural phenotyping system for at least one week where a stable food intake was achieved. Mice did not receive any interventions. Two-way ANOVA for assessment of main diet effect, virus effect and/or interaction between diet and virus effect, followed by the Sidak post-hoc test. Dark phase: food intake, diet**, $p=0.003$; water intake, diet*, $p=0.01$; energy expenditure, diet****, $p<0.0001$; average RER, diet****, $p<0.0001$; meal size, diet*, $p=0.006$; meal duration, diet***, $p=0.0003$. Light phase: energy expenditure, diet****, $p<0.0001$; diet***, $p=0.0003$.

### Jiang et al, Suppl Figure 6

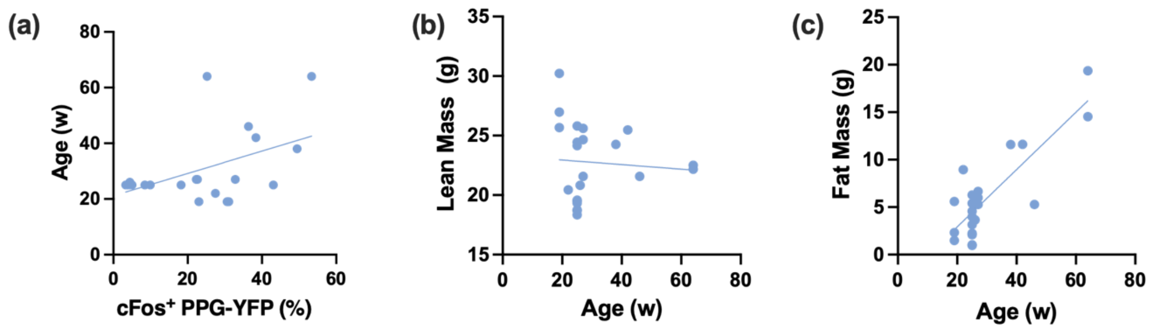


**(d)**

| Variables | Coefficient (β) | Standard Error | 95% CI | p Value |
|---|---|---|---|---|
| Intercept | -51.57 | 15.35 | (-83.82, -19.33) | **p=0.0035 |
| Lean Mass (g) | 2.68 | 0.60 | (1.43, 3.94) | ***p=0.0003 |
| Fat Mass (g) | 2.15 | 0.73 | (0.617, 3.68) | **p=0.0086 |
| Age (week) | 0.01 | 0.27 | (-0.55, 0.56) | p=0.9795 |

**Suppl Fig 6.** Positive correlation between age and $cFos^+$ PPG-YFP (%) disappears after controlling for fat mass. **a-c** Simple linear regression revealed that age is positively correlated with the percentage $cFos^+$ PPG neurons positively correlated with (slope = 0.40 ± 0.16, p = 0.02, R2 = 0.24) and fat mass (slope = 0.30 ± 0.045, p <0.0001, R2 = 0.69). **d** Multiple linear regression including age, lean mass, and fat mass revealed significant positive associations with lean and fat mass, but not with age ($R^2$ = 0.76). *p < 0.05, **p < 0.01, ***p < 0.001, ****p < 0.0001.

Jiang et al, Suppl Figure 7

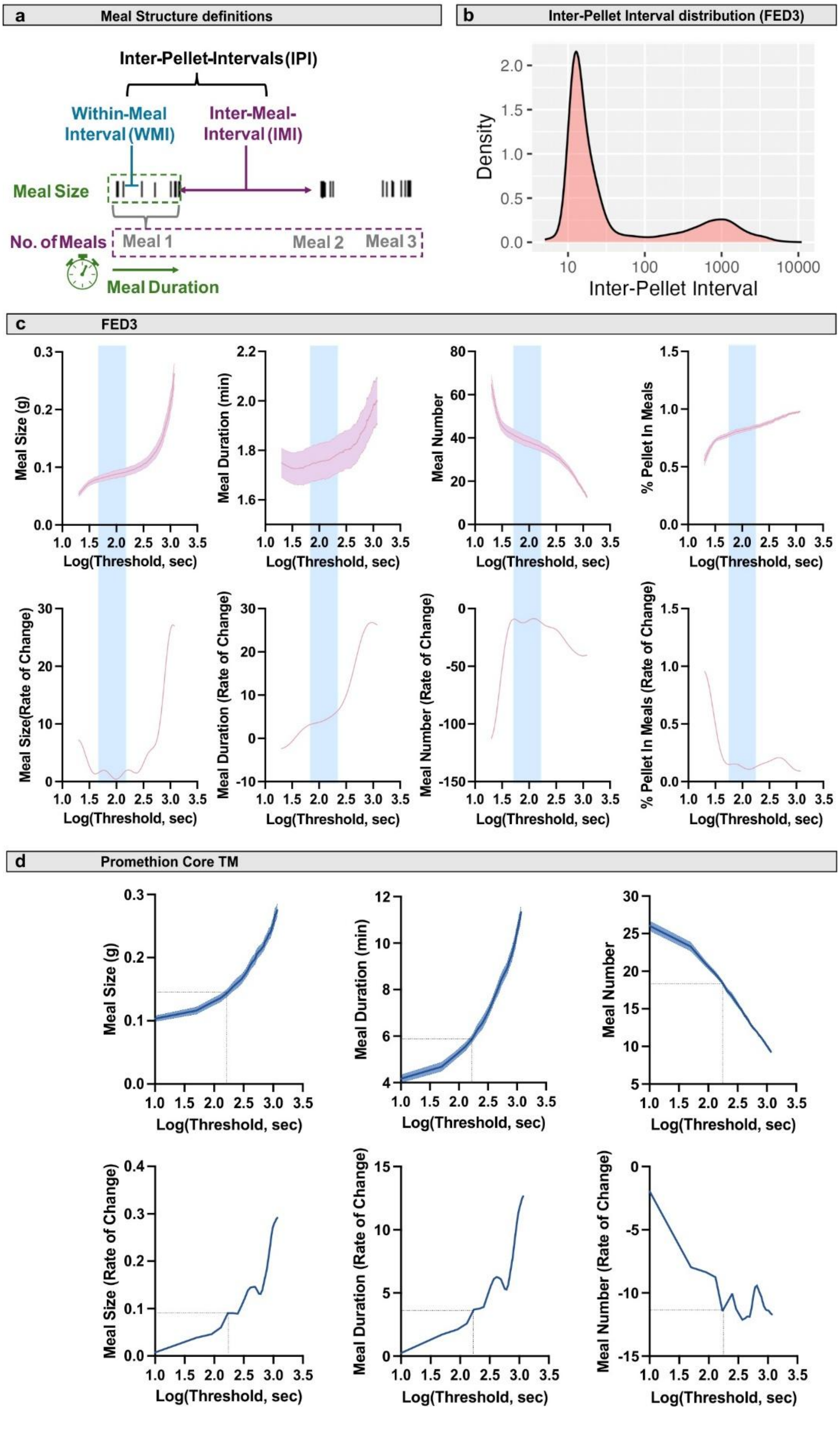


**Suppl Fig 7. Generation of meal definitions. a** Illustration of meal structures and temporal patterns, with definitions adapted from Watts et al. (2022). FED3 pellets are shown as individual bars, with a meal defined as a cluster of eating bouts separated by inter-meal intervals (IMI). Meal size refers to the amount of food consumed within a meal, while meal duration is the time taken to consume it. Satiation is assessed through meal size and duration, while satiety is evaluated using IMI and meal frequency (the number of meals consumed within a specific period). **b** Distribution of all the inter-pellet intervals (IPIs) from *ad libitum*-fed mice (n=13) across 3 experimental days. **c** Feeding data from the FED3 system were used to calculate meal size, duration, meal number, and percentage of pellets classified into meals across a range of tested thresholds. The area with minimal rate of change in the curves is highlighted in grey. Data presented as mean ± SEM, with SEM indicated by the shaded area. **d** Feeding data from the Promethion Core system (n=16), with the selected threshold of 2.24 indicated by a dashed line. This threshold corresponds to the time just before a significant change in the rate of change is observed.
